# Nimodipine prevents the development of spasticity after spinal cord injury

**DOI:** 10.1101/639211

**Authors:** Maite Marcantoni, Andrea Fuchs, Peter Löw, Ole Kiehn, Carmelo Bellardita

## Abstract

Spasticity, one of the most frequent comorbidities of spinal cord injury (SCI), disrupts motor recovery and quality of life. Despite major progress in neurorehabilitative and pharmacological approaches, no curative treatment for spasticity exists. Here, we show in a mouse model of chronic SCI that treatment with nimodipine — an FDA-approved L-type calcium channel blocker — starting in the acute phase of SCI completely prevents the development of spasticity measured as increased muscle tone and spontaneous spasms. The aberrant muscle activities are permanently blocked even after termination of the treatment. Constitutive and conditional silencing in neuronal subtypes of Ca_*V*_ 1.3 channels shows that preventive effect of nimodipine on spasticity after SCI is mediated by the neuronal Ca_*V*_ 1.3 channels. This study identifies a potentially curative treatment protocol with a specific target for the prevention of spasticity after SCI.

## Introduction

Spinal cord injury (SCI) in humans is a traumatic life-changing event that results in permanent sensory-motor disabilities. Traumatic SCI is characterized by an acute mechanical injury, resulting in the depression of spinal reflexes and absence of movements (1, 2). The initial insult is followed by a series of secondary damage to the injury epicenter and initially spared distal regions (1). Several months after the lesion, a significant number of SCI patients develops spasticity (70%)(3), a maladaptive state of hyper-excitability of the spinal circuitries resulting in the appearance of tonic muscle contractions and episodic muscle spasms (4, 5). Spasticity impairs residual motor function, negatively impacts the quality of life even leading to early mortality (6). Therefore, spasticity is a significant problem for people with SCI (3). Despite major progress in neuro-rehabilitative and therapeutic approaches, no curative treatments exist. Thus far, neurorehabilitation facilitates moderate motor recovery but is not effective in alleviating the hyperexcitability of the spinal circuitries (7, 8). In contrast, the main pharmacological strategies not only suppress the hyperexcitability but also any remaining motor responses (9, 10).

L-type calcium currents represent a major component of the persistent inward currents (PICs) underlying plateau potentials, a membrane property that allows neurons to sustain firing with little or no synaptic inputs (11–13). The expression of plateau potentials is enhanced in spinal neurons during the chronic phase of SCI with a direct correlation between expression of plateau potentials and appearance of spasticity (12, 14–16). In addition to a role in membrane excitability, L-type voltage-gated calcium channels are involved in calcium signaling and calcium-induced gene expression in neurons (17). Intracellular calcium concentration increases after SCI and remains high for days, activating downstream signaling networks that alter gene expression (18–21). Induction of these processes can lead to neuronal death, which can exacerbate the primary injury and result in long-lasting changes to network function (22, 23).

Since L-type calcium channels may be implicated in the appearing of spasticity after SCI, we evaluated the therapeutic potentials of nimodipine, an L-type calcium channel antagonist and FDA-approved compound. We demonstrate that nimodipine administration initiated early after SCI and for prolonged time is capable of preventing the development of spasticity in a mouse model of SCI. Utilizing transgenic mice we provide a detailed expression profile of Ca_*V*_ 1.3 channels in different neuronal and non-neuronal elements of the spinal cord and show that constitutive knockout or deletion of Ca_*V*_ 1.3 channels in neurons is sufficient to prevent the development of increased muscle tone and spasms in the chronic phase of SCI. The study provides compelling support for nimodipine as a potential strategy for treating spasticity in human after SCI.

## Results

### Nimodipine abolishes tonic muscle contraction and muscle spasms after SCI

To assess the role of L-type calcium channels in spasticity after SCI, we used a mouse model of SCI with a complete transection of the second sacral segment (S2) of the spinal cord that replicates clinical signs of human spasticity (15, 24,74–76). In this model of chronic SCI, lesioned wild-type (WT) mice soon after SCI exhibited a tail posture similar in shape to that seen in un-lesioned mice (Fig. S1A). However, an abnormal change in the tail posture gradually appeared, peaking six weeks after SCI (chronic state, Fig. S1B) and remaining so thereafter (Fig. S1) (15, 24–27). Electromyography recordings from tail muscles in lesioned WT mice revealed an increase in tonic muscle activity at the level of the curvatures that well correlated to the appearance of the abnormal tail posture (Unpaired t Test, p<0.001, Fig. S1D). We described the extent of the tonic muscle contraction by a *severity index* which uses tail bending angles and length as a proxy for changes in muscle tone level (Fig. S1D-E, see also materials and methods). In addition to the tonic muscle contraction, spasticity was characterized by spontaneous recurrent episodic muscle spasms with large increases in muscle activity (Fig. S1C)(15, 24). Small- and large-sized motor units were recruited simultaneously at the beginning of the spasms, inducing an abrupt and temporary increase in tail bending (Fig. S1C). Conversely, spasm termination appeared as decreased motor unit activity with successive muscle relaxation (Fig. S1C). Since the increase in muscle tone and muscle spasms duration provide quantitative measurements of spasticity and they reflect clinical signs of spasticity after SCI, we use these two parameters to evaluate the degree of spasticity in our study.

To evaluate the therapeutic action of blocking L-type calcium channels after SCI and its translational potential in humans, we administered subcutaneously nimodipine (10 mg/kg). Nimodipine is an L-type calcium blocker used for treatment of cardiovascular diseases as well as a protective agent for preventing worsening of neurological disorders after stroke (28). The drug efficiently passes the blood brain barrier allowing systemic delivery with no major side effects (29, 30). Three groups of lesioned mice were used for this study. A first group of WT lesioned mice received just vehicle solution soon after SCI (vehicle group, Fig. 1A). A second group received nimodipine from one day post-SCI (early treated) and a third group 6 weeks after SCI when spasticity is already developed (delayed treated group, Fig. 1A). The severity index and the spasm duration for all groups were assessed on the last day of treatment. Vehicle-treated animals showed strong tail bending (0.8 ± 0.05) and sustained spasms duration (0.56 ± 0.03) comparable to untreated WT lesioned animals (Unpaired t Test, p>0.05). In stark contrast, the early treated group exhibited a drastic reduction of the severity index (0.11 ± 0.02) and the muscle spasm duration (0.1 ± 0.03) when compared to the vehicle treated group (Fig. 1B-C). Similarly, the delayed treated group exhibited a significant reduction of the severity index (0.42 ± 0.04) and muscle spasm duration (0.28 ± 0.06) when compared to the vehicle-treated animals (Fig. 1B-C). However, this reduction was significantly less pronounced than that observed in the early treated animals.

**Fig. 1.**
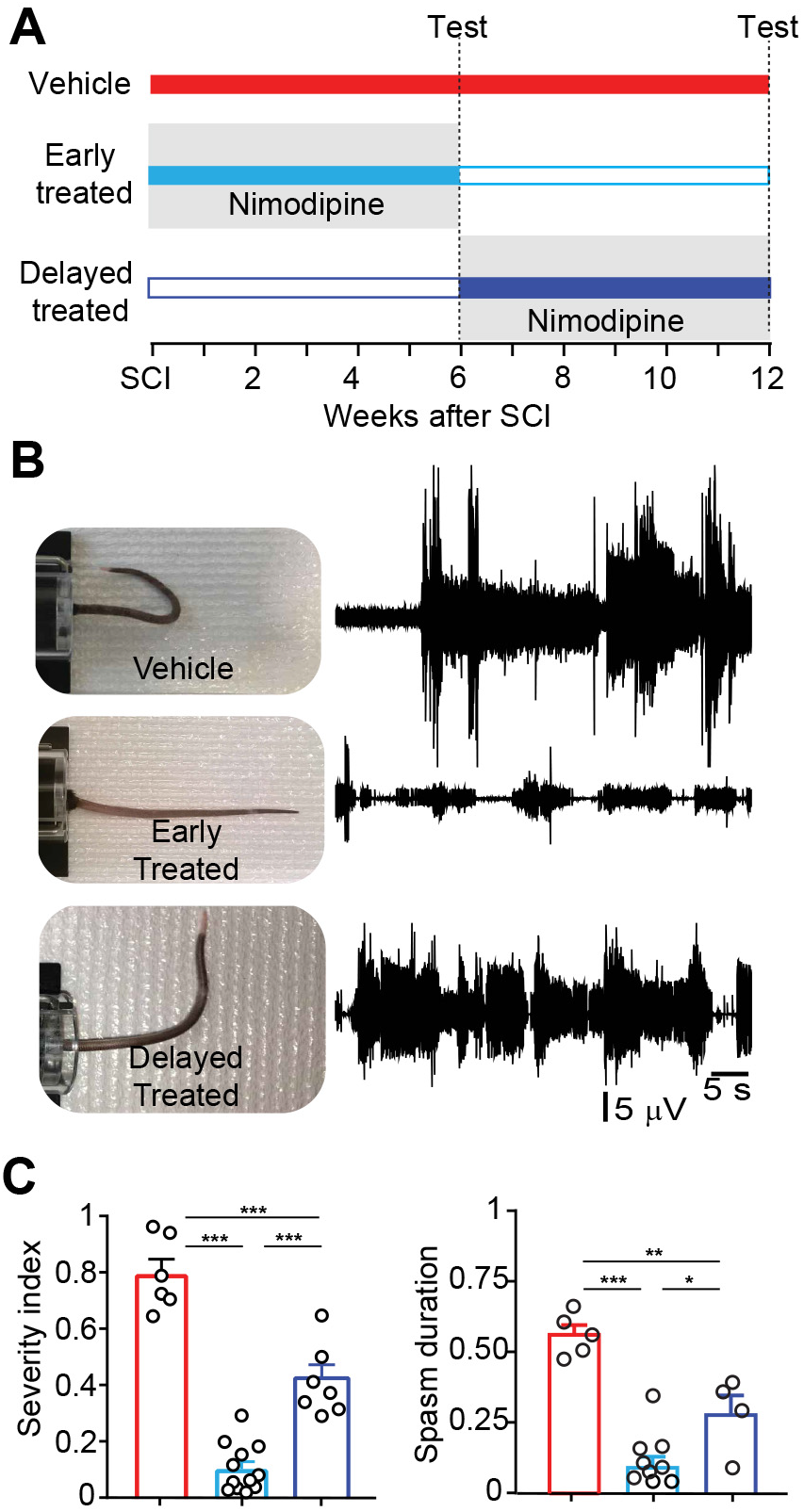
Treatment with nimodipine abolishes tonic muscle contraction and muscle spasms after SCI. **A.**Experimental protocol. Tonic muscle contraction and muscle spasms were measured at 1 week, 6 weeks and 12 weeks after SCI. Nimodipine treatment (10 mg/kg) started the day after SCI for the early treated mice and 6 weeks after SCI for the delayed treated mice. The treatment lasted 6 weeks for all groups. **B.** Representative images of mouse tails and EMG recordings from a proximal ventral muscle of the tail in vehicle-treated (upper trace), early-treated (middle trace) and delayed-treated (lower trace) mice 6 weeks after treatment. **C.** Severity index and spasms duration in vehicle-treated (red), early-treated (cyan) and late nimodipine-treated mice (blue). Each dot represents one animal. Mean ± SEM, one-way ANOVA, Tukey’s post hoc, *p<0.05, **p<0.01, ***p<0.001.

Taken together, the data indicates that prolonged treatment with nimodipine after SCI efficiently reduces the tonic muscle contraction and muscle spasms occurring after SCI. The significant albeit reduced effect of nimodipine in chronically-injured animals (delayed treated) suggests that tonic muscle contraction and muscle spasms are directly supported by calcium influx through the L-type calcium channels. The stronger effect of the early onset treatment that resulted in an almost complete abolishment of the tonic muscle contraction and spasms indicates that the early administration of nimodipine reinforce its action by an additional effect on the network than just directly blocking the L-type calcium currents.

### Only early and prolonged delivery of nimodipine prevents development of tonic muscle contraction and muscle spasms after SCI

To provide further evidence for this hypothesis, we took advantage of the possibility that the adult sacral spinal cord can be isolated six week after SCI when spasticity is fully developed (15, 24). In this way, we can study the spinal activity and the motor responses at the cellular level in animals that either received an early nimodipine treatment or none at all. To be able to visualize interneuronal activity we performed the lesions in *Vglut2*^*Cre*^;*GCaMP3*’ mice. Activity of the motor neurons was then recorded in ventral roots during simultaneous imaging of spinal interneuron activity during sensory-induced spasms (Fig. 2A, See (15)). In spinal cords from non-treated animals sustained activity was elicited by sensory stimulation in excitatory spinal interneurons resulting in prolonged motor responses (black traces in Fig. 2B). Subsequent application of nimodipine resulted in a significant reduction of interneuronal activity (mean reduction 45 ± 3.2%) and motor responses (main reduction 53 ± 5.6%, orange traces in Fig. 2B-C). This reduction induced by acute administration of nimodipine *in vitro* was similar to what was seen *in vivo* in the delayed treated animals (main reduction in spasms duration compared to the vehicle group, 49 ± 5.2%). In contrast, in spinal cords from early treated mice the duration and amplitude of the motor activity evoked by sensory stimulation was already shorter than in the non-treated animals (mean reduction 87 ± 11%, Fig 2D,) with little further reduction induced by acute nimodipine application. This result is also reminiscent of the spasms reduction observed *in vivo* in the early treated group (compared to the vehicle group 85 ± 4.1 and 81 ± 2.3% respectively). The similarity between the *in vivo* and *in vitro* results suggested that nimodipine may have a protective effect on the spinal circuitries only when given soon after SCI that actually may persist after the end of the treatment.

**Fig. 2.**
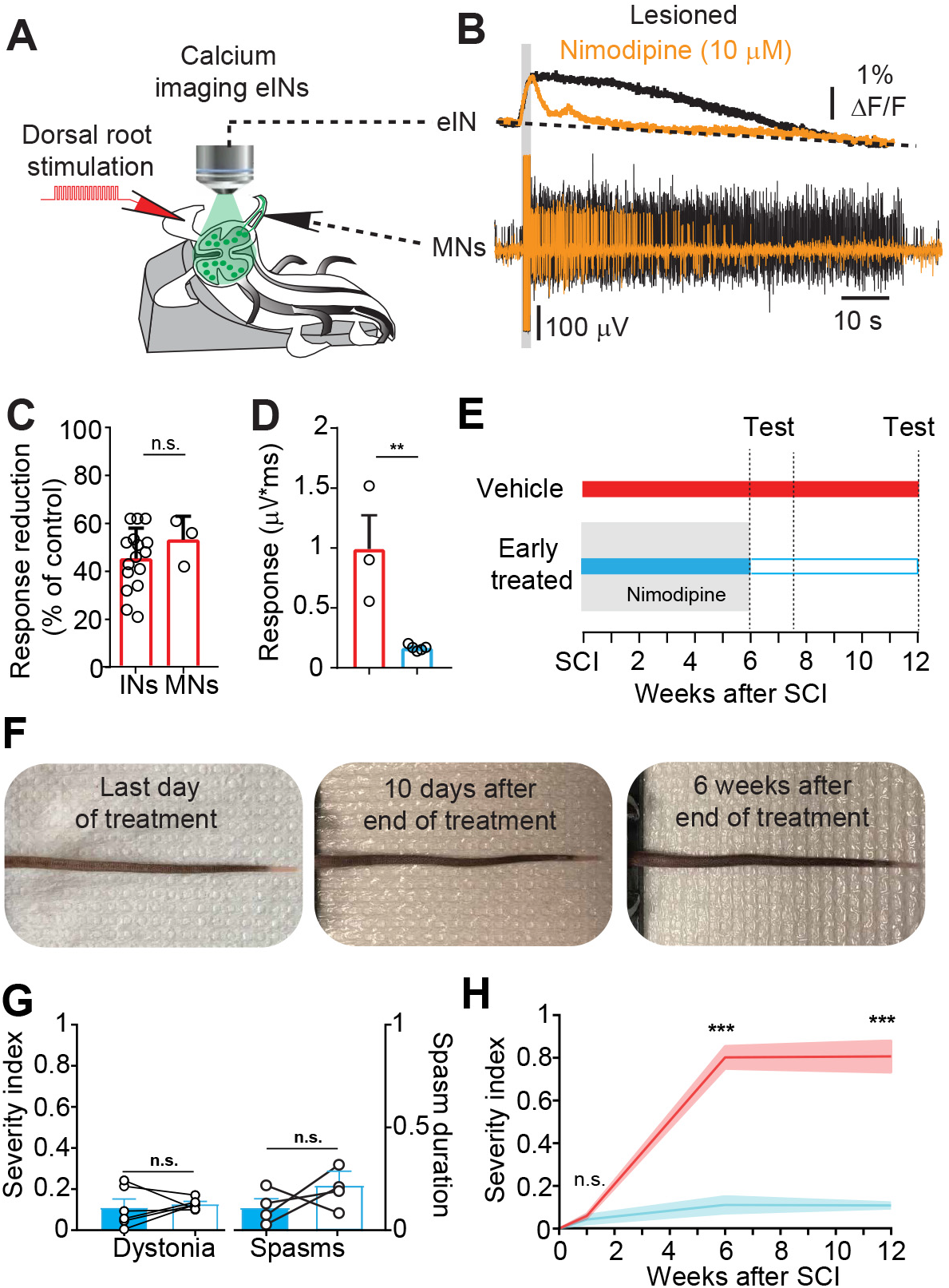
Early and prolonged treatment with nimodipine prevents tonic muscle contraction and muscle spasms after SCI. **A.** Schematic of the sacral spinal cord preparation from a lesioned *Vglut2*^*Cre*^;*GCaMP3*’ mouse 6 weeks after SCI during ventral root recording, dorsal root stimulation (stimuli of 50 *µ*s, 10 Hz, 15 *µ*A) and simultaneous imaging of the calcium dynamics of excitatory spinal interneurons. **B.** Representative ventral root recording and calcium dynamic of an excitatory interneuron (∆F/F) in a lesioned mouse during dorsal root stimulation before and after bath application of Nimodipine. **C.** Reduction of calcium responses for the spinal interneurons (n=16) and motor neurons (n=9) in three lesioned mice after nimodipine application. Mean ± SEM, unpaired Student’s t test, n.s. p>0.05. **D.** Motor response duration in lesioned non-treated (N=3, Red), lesioned and early-treated (N=5, Cyan) 6 weeks after SCI (mean ± SEM, unpaired Student’s t test, **p<0.01). **E.** Schematic of the time points (test) showed in the next panels. **F.** Representative images of the tail of an early-treated mouse the last day of treatment, 10 days and 6 weeks after termination of treatment. **G.** Severity index (left, n=5) and spasm duration (right, n=4) in early nimodipine-treated mice at the last day of treatment (blue bar) and 10 days after the end of the treatment (empty bar). Mean ± SEM. Paired t test, n.s. p>0.05. **H.** Severity index in vehicle-treated (red, n=6) and early-treated mice (cyan, n=6) over the entire experimental protocol (Mean ± SEM, paler lines indicates SEM, Two-way Anova, Sidak’s multiple comparisons.***p<0.001).

### The effect of Nimodipine persists after termination of drug treatment

To examine whether a protective effect of the nimodipine-treatment on spasticity exists, and if so, to which extent it contributes to a real preventive action on the development of spasticity, we measured tonic muscle contraction and muscle spasms in lesioned mice after the termination of the treatment in the early treated group. 10 days after the end of treatment, a time where the drug is completely cleared from the body(30) (Fig. 2E-F), there was no significant difference in the severity index and the muscle spasm duration compared to the last day of the treatment (Fig. 2G). This indicates that early nimodipine-treatment prevents the development of tonic muscle contraction and muscle spasms with a long-lasting effect, even in the absence of the drug. Even more striking, this protection appeared to be permanent. Six weeks after the termination of the drug treatment – the same time span usually required to fully develop motor dysfunctions after SCI in the absence of drug –there was no significant increase of tonic muscle contraction (Fig. 2F and 2H). Taken together, these results indicate that early and prolonged blockade of L-type calcium channels soon after SCI permanently prevents the development of increased tonic muscle activity and muscle spasms, typical of the chronic phase of SCI. It seems that nimodipine may have a dual action in preventing spasticity after SCI: a direct effect mediated by blocking the L-type calcium channels and an indirect effect that can be exerted only when the drug is given soon after the injury and permanently prevents the hyper-excitability of spinal cord circuitries below the injury.

### Genetic silencing of L-type calcium channels Ca_*V*_ 1.3 abolishes tonic muscle contraction and muscle spasms after SCI

Next, we investigated which of the L-type calcium channels nimodipine is blocking to prevent spasticity. Since L-Type calcium channels cannot be pharmacologically distinguished (12, 16, 31, 32), we used a genetic approach to answer the question. Since plateau potentials are triggered and driven by strong dendritic calcium current we focus our study on the L-type calcium channels Ca_*V*_ 1.3 that are mainly located in the dendritic branches of spinal neurons (17). We therefore used a *Cacna1d* knockout mouse that exhibits functional silencing of the Ca_*V*_ 1.3 calcium channels (33) (Ca_*V*_ 1.3 KO, Fig. S2 and Fig. 2A). In contrast to the lesioned WT previously described (Fig 3B, Fig S1), the tail of chronically lesioned Ca_*V*_ 1.3 KO mice did not show chronic bending with the gross tail appearance remarkably similar to un-lesioned WT mice and early treated mice (Fig. 3B and Fig. S3A). Electromyography on the tail muscles showed little tonic activity and sporadic small amplitude spasms (Fig. 3C). In chronically-injured WT mice this severity index ranged from 0.65 to 0.95 (with a mean value of 0.81 ± 0.03– with 1.0 being the most severe case, Fig. 3D-E and Fig. S1E-F). In contrast, chronically-lesioned Ca_*V*_ 1.3 KO mice exhibited a severity index close to zero (mean value of 0.05 ± 0.005, Fig. 3D-E; Fig. S3D-E) with tonic activity not significantly different from un-lesioned WT mice and much lower than chronically-lesioned WT mice (Fig. S3C). The severity index of the lesioned WT and Ca_*V*_ 1.3 KO mice remained the same for the twelve weeks period of observation (Fig. 3G). Simultaneously, lesioned Ca_*V*_ 1.3 KO mice exhibited a strong reduction in muscle spasm duration compared to chronically lesioned WT mice (Fig. 3C and 3F), with favored recruitment of smaller-sized motor units (Fig. S3B). Together, these results demonstrate that blockage of Ca_*V*_ 1.3 channels is sufficient to prevent spasticity after SCI. These results also indicate that the preventive effect of the nimodipine treatment after SCI is mainly mediated by the blockage of the Ca_*V*_ 1.3 channels.

**Fig. 3.**
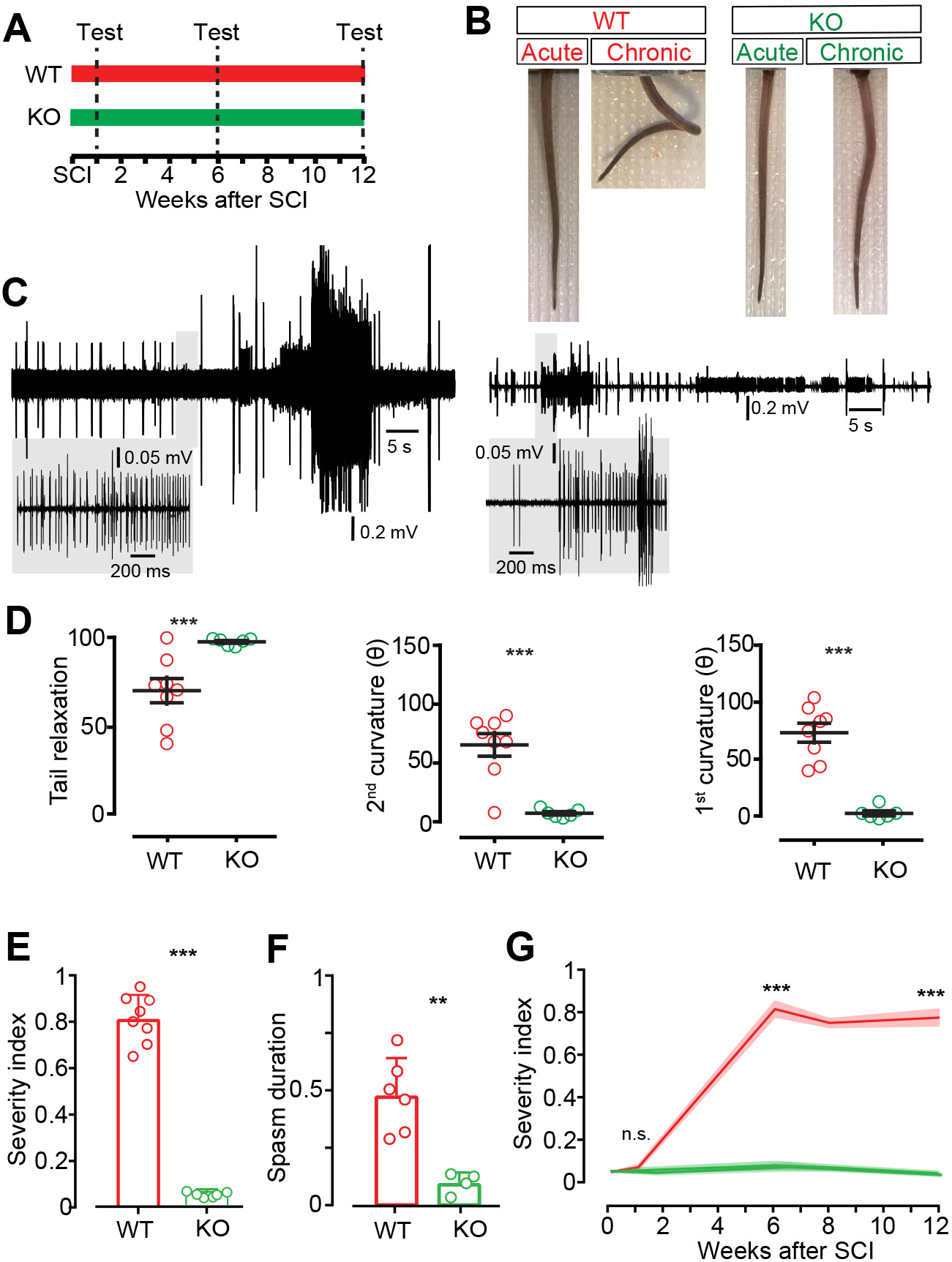
Blockade of L-type calcium channels Ca_*V*_ 1.3 is sufficient to prevent tonic muscle contraction and muscle spasms after SCI. **A.** Schematic of the experimental protocol. **B-C.** Representative images (B) and electromyography on the ventral muscle of the tail (C) in wild-type mice (left) and Ca_*V*_ 1.3 KO mice (right) the day after (acute) and 6 weeks after (chronic) SCI. **D.** Tail relaxation and angles of the first and second curvature for wild-type (red, WT, n=8) and Ca_*V*_ 1.3 KO (green, KO, n=6) mice 6 weeks after SCI (mean ± SEM, unpaired Student’s t test, ***p<0.001). **E-F.** Severity index (E, mean ± SEM, unpaired Student’s t test, ***p<0.001) and spasm duration (F, mean ± SEM, unpaired Student’s t test, ***p<0.001) 6 weeks after SCI. **G.** Mean severity index at 0, 1, 6 and 12 weeks after SCI in lesioned wild-type (red, n=8) and lesioned Ca_*V*_ 1.3 KO mice (green, n=6). Paler colors indicate SEM. Two-way Anova, Sidak’s multiple comparisons.

### Ca_*V*_ 1.3 calcium channels are abundantly expressed in neurons and non-neuronal elements of the spinal cord

Next, we examined the spatial distribution of Ca_*V*_ 1.3 channel expressing cells in the mouse spinal cord to identify which cell populations may be involved in the development of spasticity. For this we used a Ca_*V*_ 1.3 reporter allele that, upon Cre-mediated recombination, expresses eGFP from the Ca_*V*_ 1.3 locus (*Cacna1d-eGFP*^*flex*^, Fig.S2B and(34)). We crossed *HoxB8*^*Cre*^ mice (35) with the *Cacna1d-eGFP*^*flex*^ reporter to obtain reporter expression in cells caudal to the 4^*th*^ cervical segment (Fig. S2C). Ca_*V*_ 1.3 calcium channels were widely expressed in the spinal cord from early development (Fig. S4A-C) to adulthood (Fig.4). Approximately 55% (51 ± 2.1, mean ± SEM) of the spinal cells expressed Ca_*V*_ 1.3 channels (Ca_*V*_ 1.3^+^ cells, Fig 4A-B). Ca_*V*_ 1.3^+^ cells were found along the rostrocaudal axis of the cord (Fig. 4C-D) distributed across the dorsal (laminae I-V), intermediate (laminae VII-VII and X) and ventral (laminae VIII-IX) laminae (Fig. 4E-H). Ca_*V*_ 1.3^+^ cells exhibited the highest density in the superficial dorsal horn in all segments (Fig. 4H) with a lower density along the dorsoventral and mediolateral axes of the cord (Fig. 4E-H), following the typical neuronal distribution within the grey matter of the spinal cord (36).

**Fig. 4.**
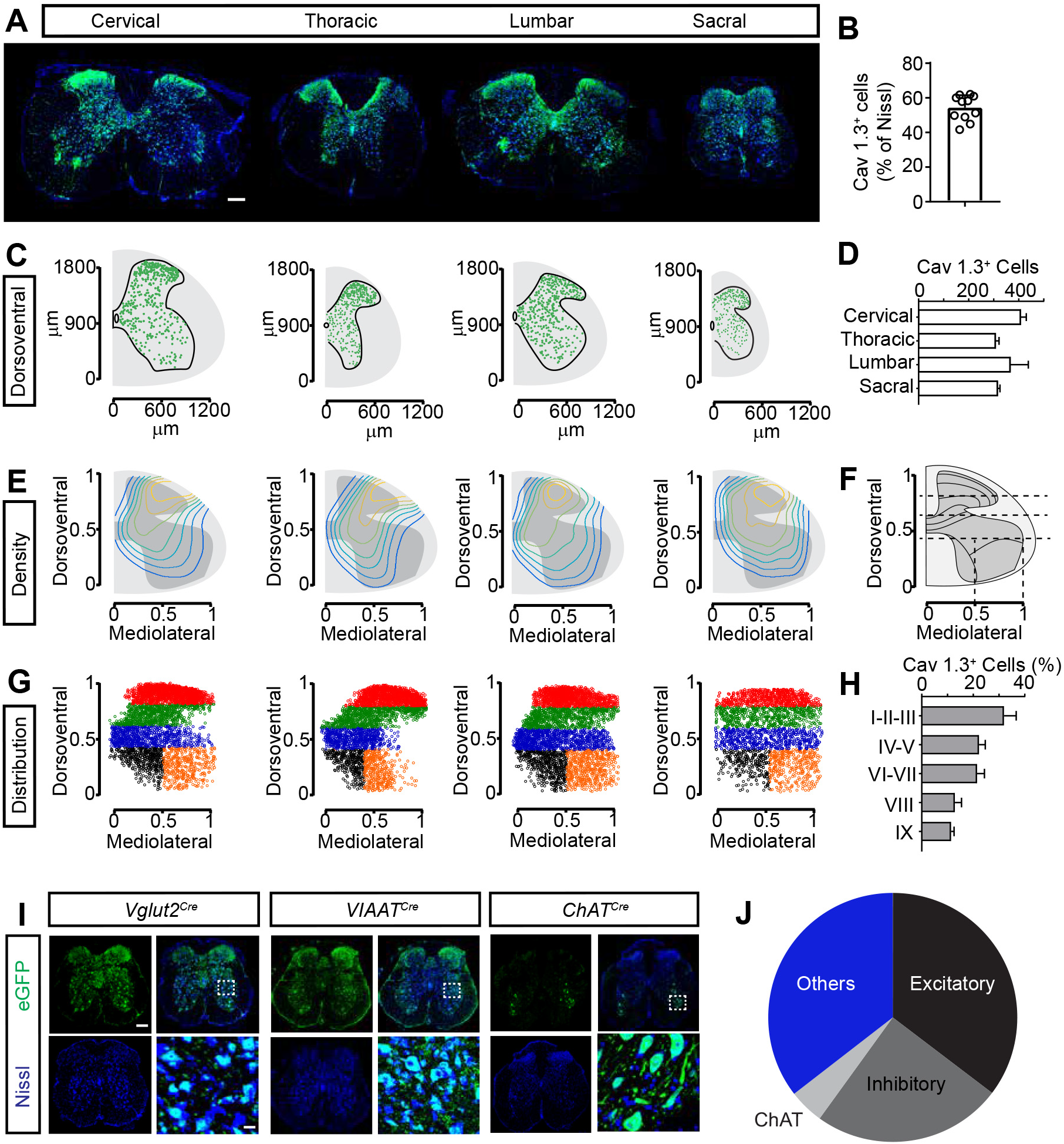
Ca_*V*_ 1.3 channels are abundantly expressed in the mouse spinal cord. **A.** Representative transverse sections (maximal projections) of the cervical, thoracic, lumbar and sacral spinal cord in *HoxB8*^*Cre*^;*Cacna1d-eGFP*^*flex/+*^ adult mice. Scale bar = 200 *µ*m. **B.** Average number of cells expressing Ca_*V*_ 1.3 channels (GFP, green) out of all spinal cells (Nissl) in the same section (n=12, N=3). **C.** Digital coordinate of Ca_*V*_ 1.3 cells from the left spinal cords in A. **D.** Grand mean of Ca_*V*_ 1.3^+^ cells in different segments of the spinal cord (data from B). **E.** Laminar density distribution of Ca_*V*_ 1.3^+^ cells with the highest density in yellow and lowest density in blue. **F-H.** Digital normalization of the Ca_*V*_ 1.3^+^ cell position (F) and subsequent clustering of Ca_*V*_ 1.3^+^ positions in laminae (G-H) (cervical = 3657, thoracic = 3406, lumbar = 4114, sacral = 3029). **I.** Transverse sections (maximal projections) of the sacral spinal cord from adult *Vglut2*^*Cre*^;*Cacna1d-eGFP*^*flex/+*^, V*IAAT*^*Cre*^;*Cacna1d-eGFP*^*flex/+*^ and *ChAT*^Cre^;*Cacna1deGFP*^*flex/+*^ mice stained for eGFP (green) and Nissl (blue), merged and magnified (dotted square). Scale bar = 200 *µ*m (20 *µ*m in the magnified area). **J.** Proportion of excitatory neurons (*Vglut2*^+^, N=2), inhibitory neurons (*VIAAT*^+^, N=2), motor neurons (*ChAT*^+^, N=2), and non-neuronal cells (others) expressing Ca_*V*_ 1.3 channels (% of the Nissl stained cells)

To assess the expression of the Ca_*V*_ 1.3 channels in different neural populations we crossed the *Cacna1d-eGFPflex* with *Vglut2*^*Cre*^(37), *VIAAT*^*Cre*^(38) and *ChAT*^*Cre*^(39) mice leading to reporter expression in glutamatergic, GABAergic/glycinergic, or cholinergic neurons, respectively. By comparison with the *HoxB8*^*Cre*^, we found that most of the spinal Ca_*V*_ 1.3^+^ cells were of neuronal origin with 35% (± 6.5) of them being *Vglut2*^+^, 25% (± 5) being *VIAAT*^+^ and 5% (± 2.1) *ChAT*^+^ (Fig. 4I-J). Almost all *ChAT*^+^ ventral horn motor neurons (somatic motor neurons, S-MNs) and lateral horn preganglionic motor neurons (P-MNs) expressed Ca_*V*_ 1.3 channels whereas only a small proportion of cholinergic interneurons were Ca_*V*_ 1.3^+^cells (Fig. S4D-H).

In conclusion, Ca_*V*_ 1.3 channels are abundantly expressed in the spinal cord, including neuronal elements of each neurotransmitter-defined subtypes (65 ± 3.4%) and non-neuronal elements (35 ± 9.8%).

### Neuronal Ca_*V*_ 1.3 channels is directly involved in the generation of tonic muscle contraction and muscle spasms

Given the abundant expression of Ca_*V*_ 1.3 channels in neuronal and non-neuronal elements of the spinal cord, we wondered whether the preventive effect of silencing the Ca_*V*_ 1.3 channels on spasticity after SCI may be of neuronal origins. We have previously shown that both excitatory (*Vglut2*^+^) and inhibitory (*VIAAT*^+^) interneurons are recruited during spasms (15) and a large proportion of these neuronal populations expresses Ca_*V*_ 1.3 channels (Fig. 4). We therefore generated *Vglut2*^*Cre*^; Ca_*V*_ 1.3KO and *VIAAT*^*Cre*^; Ca_*V*_ 1.3KO mice (Fig. 5A).

**Fig. 5.**
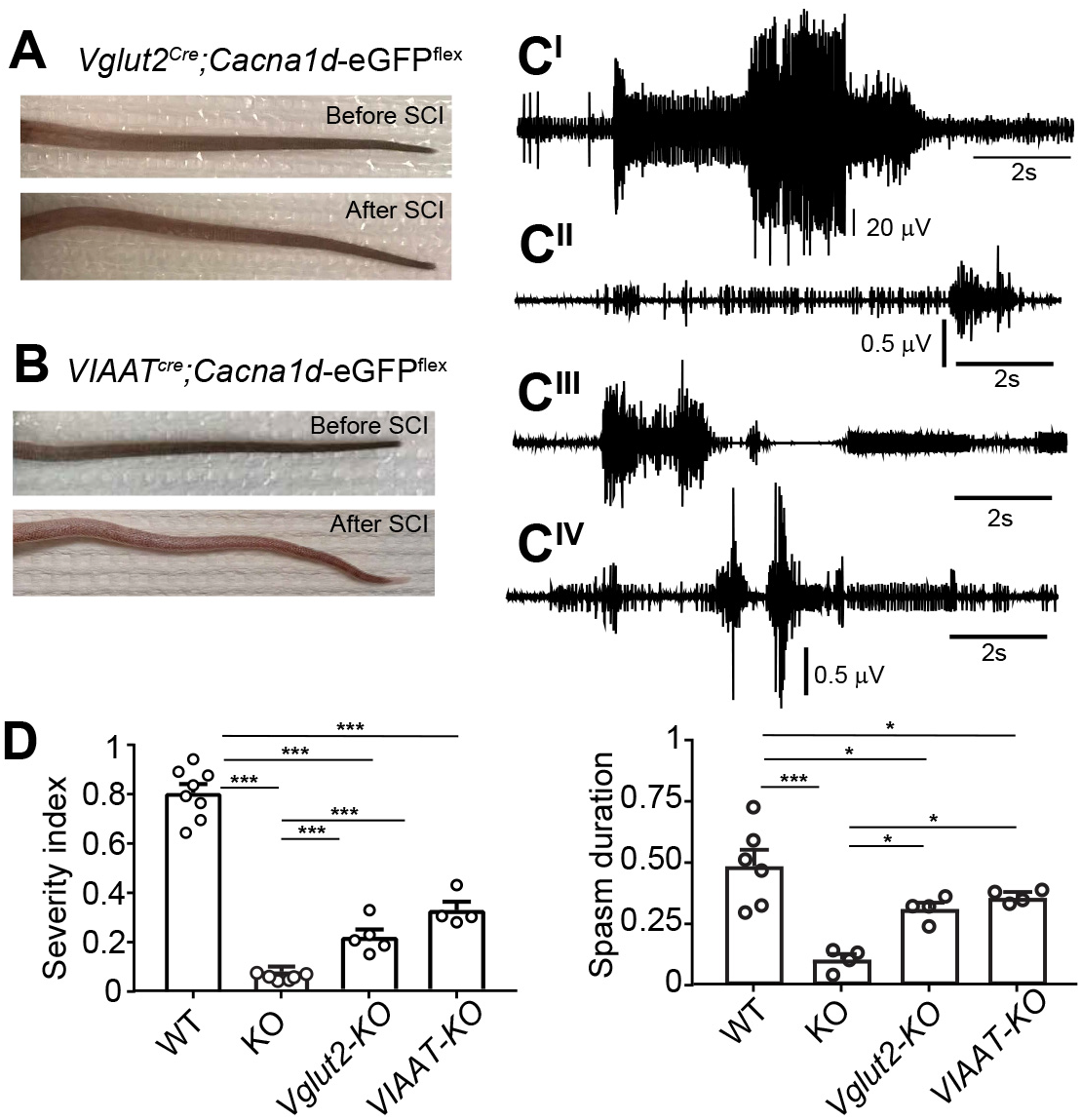
Involvement of the neuronal Ca_*V*_ 1.3 channels in the appearance of tonic muscle contraction and muscle spasms after SCI. **A-B.** Images of mouse tails with genetic silencing of the Ca_*V*_ 1.3 calcium channels in excitatory (A, *Vglut2*^*Cre*^;*Cacna1d-eGFP*^−/−^) and inhibitory (B, *VIAAT*^*Cre*^; *Cacna1d-eGFPtextsuperscript*-/-) neurons before and six weeks after SCI. **C.** Representative EMG recordings from ventral muscles of the mouse tail in wild-type (CI), Ca_*V*_ 1.3 KO (CII), *Vglut2*^*Cre*^;*Cacna1d-eGFP*^−/−^ (CIII) and *VIAAT*^*Cre*^; *Cacna1d-eGFP*^−/−^ (CIV) mice 6 weeks after SCI. **D.** Quantification of *severity index* (right) and spasms duration (left) 6 weeks after SCI in wild-type (WT),Ca_*V*_ 1.3 KO (KO), *Vglut2*^*Cre*^;*Cacna1d-eGFP*^−/−^ (Vglut2-KO) and *VIAAT*^*Cre*^; *Cacna1d-eGFP*^−/−^ (VIAAT-KO) mice. Each dot indicates an animal. Mean ± SEM, Ordinary one-way Anova, Holm-Sidak’s multiple comparisons test, with a single pooled variance. *p<0.05, **p<0.01, ***p<0.001.

To determine the specificity of the KO allele, we measured persistent inward currents (PICs) in spinal neurons (40–42) from newborn mice with one functional copy of the channel (Ca_*V*_ 1.3^+/−^) or with the complete silencing of the Ca_*V*_ 1.3 gene (Ca_*V*_ 1.3^−/−^) in spinal neurons (Fig. S5). The persistent sodium-dependent component was blocked by TTX, leaving the persistent calcium-dependent and TTX-resistant PICs (40). About half of the PICs (52 ± 10%) were generated by L-type calcium currents (nifedipine-sensitive) in Ca_*V*_ 1.3^+/−^ while the nifedipine-sensitive component in the Ca_*V*_ 1.3^−/−^ was greatly reduced (19 ± 2.9%, Fig. S5A-D). The I-V curves of the nifedipine-sensitive current displayed similar activation and peak voltages but its intensity was drastically reduced in Ca_*V*_ 1.3^−/−^ (Fig. S5C-E). When measured by subtraction, Ca_*V*_ 1.3 calcium component represents about 35% of the PICs in spinal neurons (Fig. S5E). All together these results show that Ca_*V*_ 1.3KO allele efficiently silences the Ca_*V*_ 1.3 calcium channels in neurons and that a minor nifedipine-sensitive PIC, presumably generated by Ca_*V*_ 1.2 calcium channels, is left in the Ca_*V*_ 1.3KO mice.

Six weeks following complete transection injury, lesioned *Vglut2*^*Cre*^;Ca_*V*_ 1.3KO and *VIAAT*^*Cre*^;Ca_*V*_ 1.3KO mice exhibited a diminished bending of the tail at rest and lower muscle activity (Fig. 5B-D). Both the tonic muscle contraction and the duration of muscle spasms were significantly reduced in the two Ca_*V*_ 1.3KO mice compared to lesioned WT mice (Fig. 5E). However, the reduction of the *severity index* and the muscle spasm duration in these mice was not as drastic as in the constitutive Ca_*V*_ 1.3 KO (Fig. 5E). It is worth noting that the silencing of the channels in the excitatory neurons may additionally silence the Ca_*V*_ 1.3 channels expressed in some *Vglut2*^+^ motor neurons (43–45).

These data demonstrate that neuronal silencing of Ca_*V*_ 1.3 channels strongly reduces the tonic muscle contraction and muscle spasms after SCI. However, the more pronounced effect of abolishing spasticity after SCI in the constitutive Ca_*V*_ 1.3 KO (Figure 3) likely reflects the role of multiple subpopulations of spinal neurons in the generation and maintenance of spasticity after SCI (15) with a possible minor contribution of Ca_*V*_ 1.3^+^ non-neuronal cells (46, 47).

## Conclusions

Spasticity, meant as muscle stiffness and muscle spasms, is highly associated with severe motor incomplete cervicothoracic injuries (3, 48, 49). The present study provides a basis for a potential curative treatment of problematic spasticity after SCI. We demonstrate that nimodipine prevents the development of the aberrant motor activity with long-lasting effects on the motor symptoms - acting as a curative treatment paradigm in this mouse model of SCI. Its effect is optimal when given early after the SCI and it is mediated by blocking the activity of the neuronal Ca_*V*_ 1.3 channels. Together these findings identify a mechanism for development of spasticity and provide a potential new therapeutic strategy that can be translated in humans for the prevention of spasticity after SCI.

Baclofen represents the main antispastic medication for patients and heavily impacts any voluntary and motor responses as well as cognitive functions (50). Nimodipine is a well-tolerated drug for management of cardiovascular diseases and subarachnoid stroke with no major side effects (28). It is currently under investigation for treatment of neurological diseases including Parkinson’s and multiple sclerosis in which cell vulnerability has been largely attributable to the increased cell excitability mediated by calcium influx through Ca_*V*_ 1.3 calcium channels (46, 51). Recently, a large observational cohort study described that early and only early application of gabapentinoids, negative modulators of calcium channels, significantly improved motor recovery in SCI patients (52). Since spasticity represents a major impediment to movement, altering posture and gaits, and reduced calcium currents may promote more physiologic operation of neural circuits (16, 53), nimodipine treatment may be a useful tool when combined with other pharmacological and neurorehabilitative approaches commonly used to treat motor dysfunction after SCI. Although, we did not combine nimodipine with neurorehabilitation paradigms, it is worth to investigate their potential synergy for improving recovery. Moreover, since spasticity is commonly expressed in other neurological disorders like cerebral palsy, stroke, amyotrophic lateral sclerosis and multiple sclerosis, the therapeutic action of the drug treatment may be of interest for several neurodegenerative pathologies. Thus, the treatment protocol with nimodipine outlined here is potentially translatable to human SCI patients. However, further studies are required to deeper understand the drug effects. Our mouse model of SCI with a full transection of the sacral spinal cord only affects tail muscles. More clinical models, like cervical/thoracic contusion SCI models, should be used to verify the drug effects on movements including grasping and locomotion that are usually heavily compromised in SCI. However, it should be noticed that no other animal model of SCI express signs of spasticity (54) as well as the SCI model used in this study. Furthermore, the advantage of the model is that it allows studies of the cellular mechanisms underlying the drug effects in the *in vitro* preparation.

The effect of nimodipine on spasticity seems to be explained by a dual action in the spinal cord. A primary effect exerted independently by the time of administration after SCI and common to all groups examined on the study. Nimodipine directly blocks L-type calcium channels lowering excitability of spinal neurons and thus, directly reduces muscle activity. L-type calcium channels are known to prolong neuronal firing in physiologic condition (11, 16, 55) and their currents in spinal neurons have been related to abnormal motor responses after peripheral and central injury (12, 16, 56). Larger calcium currents may be the result of changes in voltage-dependent activation and inactivation curves of the channels to more hyperpolarized potentials, as identified in humans presenting different neurological disorders with a common expression of spasticity (57–59) or alterated upstream regulatory elements of the channels (26). Moreover, Ca_*V*_ 1.3 calcium channels may increase neuronal excitability after injury indirectly, by cooperative gaiting of clustered Ca_*V*_ 1.3 channels (60), activating calcium activated nonspecific cation channels (16) and BK channels (61) or promoting activation of calmodulin or calcium-dependent calpains to increase sodium channels activity after SCI (62, 63). In our study nimodipine treatment does not completely block the sustained motor contraction in the chronic un-treated animals indicating that other cellular mechanisms than Ca_*V*_ 1.3 channels support the the expression of spasticity. The remaining aberrant motor activity may be supported by the persistent sodium currents (12, 62). However, the contribution of the sodium currents to spasticity disappeared when nimodipine is applied soon after SCI, revealing the secondary effect of nimodipine in abolishing spasticity after SCI. This secondary effect is revealed as a specific therapeutic window that maximizes the drug effects and allows for permanent protection. Since L-type calcium channels represent a major source of calcium ions in neurons and intracellular Ca2^+^ increases soon after trauma lasting for weeks after it (21), the early treatment with nimodipine may simultaneously interfere with different calcium-dependent processes that contribute to reorganization of spinal circuitries after SCI. First, genetic silencing of the Ca_*V*_ 1.3 calcium channels indicates that nimodipine may directly act at the neuronal level. L-type calcium channels may induce influx of calcium ions that activate calcium-dependent lipases and/or proteases and inhibit mitochondrial function leading to cell death (64, 65), induce expression of genetic programs for axonal degeneration (66), fiber sprouting and synaptic rearrangements (23, 53, 66, 67). Additionally, Ca_*V*_ 1.3 calcium channels and, thus, nimodipine block may affect neuronal function by reducing vasospasm (68), increasing remyelination of neuronal axons (69, 70), enhancing microglia apoptosis and reducing astrocyte activation (46, 71), all processes that promote neuronal survival (72). Thus, the secondary effect on nimodipine may blunt secondary injury changes and enhance restorative processes preventing maladaptive plasticity of spinal circuitries typical of the chronic phase of SCI.

In conclusion, spasticity after SCI is characterized by multiple pathological events in different post-injury phases whose sequence and underlying cellular mechanisms are unknown. Here we have identified Ca_*V*_ 1.3 channels as trigger of multiple neural processes whose activity soon after injury leads to the development of spasticity after SCI. The translational potential of the early and prolonged blockade of the channels with nimodipine may represent a major advance in treating spasticity after SCI.

## Acknowledgments

The authors would like to thank Jörg Striessnig for providing us with the Ca_*V*_ 1.3 KO mouse. The Cacna1d-eGFPflex mouse was kindly donated by Hans Gerd Northwang. The authors thank Kajana Satkunendrarajah and Spyridon K. Karadimas for suggestions on the previous version of the manuscript. The authors thank the members of Ole Kiehn’s lab for inspiring discussions and comments on the work and the manuscript presented here.

## Author contributions

C.B. and O.K. devised the project, designed the studies. M.M. and C.B. performed the anatomical studies, all the in vivo experiments and analyzed the data. A.F. performed the patch-clamp experiments. P.L. contributed to the anatomical experiments. C.B., O.K. M.M. interpreted the results. C.B. and O.K. wrote the manuscript with contribution from all authors.

## Data and materials availability

Data and Materials will be made available through a material transfer agreement upon request to O.K.

## Materials and Methods

### Mice strains

Wild-type, *Vglut2*^*Cre*^(37), *VIAAT*^*Cre*^(38), *HoxB8*^*Cre*^(35), *Cacna1d-eGFP*^*flex*^ (34) and Ca_*V*_ 1.3 α1 deficient mice(33) (Ca_*V*_ 1.3 KO) of both sexes were used. Animals were genotyped before experiments. Animals were housed in rooms at 22 °C and 12:12 light-dark cycle. All animal experiments and procedures were approved by Dyreforsøgstilsynet in Denmark and the local ethics committee at University of Copenhagen. For conditional deletion of the *Cacna1d* gene from the gluta-matergic (glycinergic/GABAergic) population, *Vglut*^*Cre*^(*VIAAT*^*Cre*^) mice were crossed with a homozygous *Cacna1d-eGFP*^*Flex*^ mouse to obtain silencing of one allele. These mice were then crossed back with another homozygous *Cacna1d-eGFP*^*Flex*^ mouse line to obtain the silencing of the second allele (see also Fig. S1). Ca_*V*_ 1.3 KO mice were genotyped with the following primers: Ca_*V*__Fw 5-GGAGTTGTGTATATCTGTTAAGCC-3 and Ca_*V*_ KO_Re 5-CTCGTTCATATTCTAACTCCCTA-3 (product sizes: 2448 bp from the recombined *Cacna1d* locus and 1170 bp from the wild-type *Cacna1d* locus). Primers for the *Cacna1d-eGFP*^*flex*^were: Ca_*V*_-Re 5-GTCTCCCATCTTGCATTTCC-3 and Ca_*V*__Fw (product sizes: 400 bp from the recombined *Cacna1d* locus and 250 bp from the wild-type *Cacna1d* locus) see Fig. S1.

### Surgery, post-operative care

All surgical procedures were performed in adult mice (two to four months old) as previously described(15, 24). Briefly, the animal was deeply anesthetized with isoflurane and a vertical incision was made at the level of the second lumbar vertebral body to expose the second sacral spinal segment. Xylocaine (1%) was applied locally and the spinal cord tissue was aspirated using small glass pipette (diameter of 100 µm), paying attention not to damage the main arteries and veins around the cord. Once the lesion was completed the muscles surrounding the spinal column and the skin were closed with one 6.0 silk suture to protect the exposed spinal column. The animal was single caged and a post-surgery treatment of Buprenorphine (0.1 mg/Kg) and Carprofen (5 mg/Kg) was given subcutaneously for 2 to 5 days. After about 7 days, the animal was caged again with its initial partner. Only animals with a complete lesion of the spinal cord, visually inspected during the dissection were included in the study.

### Evaluation of tonic muscle contraction and muscle spasms after SCI

The mice were restrained in a mouse restrainer with the tail hanging, free to move. Recording electrodes for electromyography (EMG) were inserted intramuscularly in the ventral part of the tail at the level of the seventh/eighth coccygeal vertebral body (rostral recordings) and at the 14^th^ /15^th^ coccygeal vertebral body (caudal recordings). The electrodes consisted of two twisted Teflon-coated platinum/iridium wires (diameter of 125 µm, WPI, code number PTT0502). They were threaded through a 30 ½ gauge hypodermic needles before the Teflon cover was removed from the tips and the two electrodes were then inserted 1-1.5 mm apart in the tail. A single wire, prepared like the electrodes for the recordings, was used as ground electrode and inserted on the dorsal side of the tenth vertebral body. Since mice were spinalized they did not feel any pain during this procedure. The proximal ends of the recording wires were connected to a differential amplifier (Custom made). The EMG signal was sampled with 5 KHz, band-pass filtered (60-1000 Hz) and digitalized (Digidata 1440A, Molecular Device) for offline analysis using Spike2 (CED products). The *severity index* used to describe the change in tail morphology after lesion was generated using pictures of tail of the restrained mice. The tail was divided into 4 points (a, b, c, d) and thus three segments (A, B and C, Fig. S1D). Three measurements were taken: 1) the angle between the segments A and B; 2) the angle between the segments B and C; 3) the relationship between the length of all segments (length of the tail, A+B+C) and the shortest distance between the base of the tail (point a) and the tip of the tail (point d) (indicating a contraction of the tail, new segment D). Next, the angles were compared to their maximal value (120°) and normalized to the specific weight that was given to each parameter in the severity index. Since the first angle is the parameter that better captured the contraction of the tail we gave it a larger weight (0.4) than the two other measurements (0.3 maximum value). Then, the parameters were summed to generate the severity index with the greatest value (1) indicating the most severe tonic muscle contraction. The duration of muscle spasms were quantified by measuring their duration. The beginning of a spasm was determined when the firing frequency of the motor units increased above the background activity and similarly the spasms were considered terminated when the firing frequency returned to the background level. In the results section, spasm duration is normalized (spasms duration in seconds/total recording time).

### Ventral root recordings and calcium dynamics of excitatory neurons in the isolated spinal cord

The sacral spinal cord preparation was isolated as previously described (15, 24). Briefly, adult lesioned mice were anesthetized with isoflurane and a laminectomy was performed from the 11th thoracic vertebral body. The spinal cord caudal to the second sacral segment was removed and placed in a chamber with oxygenated modified artificial cerebrospinal fluid (in mM: 101 NaCl, 3.8 KCl, 18.7 MgCl2, 1.3 MgSO4, 1.2 Kh2PO4, 10 Hepes, 1 CaCl2, 25 Glucose; oxygenated in 100% O2). After removing the Dura Mater and shortening the roots, the spinal cord was placed in a perfusion chamber with normal Ringer solution (in mM: 111 NaCl, 3 KCl, 11 glucose, 25 NaHCO_3_, 1.25 MgSO_4_, 1.1 KH2PO_4_, 2.5 CaCl_2_ oxygenated in 95% O_2_ and 5% CO_2_ to obtain a pH of 7.4) and maintained at 22–24°C for an hour before any further manipulation. A train of stimuli (50 *µ*s, 10 Hz) was delivered to the dorsal root (usually the first coccygeal root, Co1) using a glass pipettes while the motor response was recorded in the homologue ventral root using a tight-fitting suction electrode. In case of simultaneous calcium imaging the sacral spinal cord preparation isolated from *Vglut2*^*Cre*^;*GCaMP3’* mice was transversally cut at the Co1 level and placed in the perfusion chamber with the dorsal and ventral horns facing the objective (40X). Illumination of the cord for excitation (470-490 nm) and visualization (520-560 nm) was obtained by a 100 W mercury lamp. Change in fluorescence was detected using a digital CMOS camera (Orca 4, Hamamatsu) at 10 frames/s and data were analyzed off-line using ImageJ. Change in fluorescence intensity over time was converted in ∆F/F = (Ft-F0)/F0 x 100 where Ft is the fluorescence at any time and F0 is the background fluorescence. Areas of interest (ROIs) were manually drawn over cell somas and the ∆F/F was calculated for each ROI.

### Measurement of persistent inward current (PICs)

Transverse spinal cord slices were used to investigate the PICs in neurons in heterozygote *HoxB8*^*Cre*^;*Cacna1d-eGFP*^+/−^ (one allele of the *Cacna1d* gene active) and homozygous *HoxB8*^*Cre*^;*Cacna1d-eGFP*^−/−^ (both alleles inactivated) mice. Spinal cords from newborn mice (P6-P8) were dissected set in 2% agar and transversally sectioned (350 m thick) using a vibratome (Microm: HM 650V) in ice-cold solution containing in mM: 101 NaCl, 3.8 KCl, 18.7 MgCl_2_, 25 glucose, 1.3 MgSO_4_, 1.2 KH_2_PO_4_, 10 HEPES, 1 CaCl_2_, and continuously oxygenated with 100% O_2_. After sectioned, the slices were incubated at room temperature in a normal Ringer’s solution containing in mM: 111 NaCl, 3 KCl, 11 glucose, 25 NaHCO_3_, 1.3 MgSO_4_, 1.1 KH_2_PO_4_, 2.5 CaCl_2_, pH 7.4, bubbled with 95% O_2_/5% CO_2_ for one hour before recording and then moved to a recording chamber perfused with the same solution at a flow rate of 4-5 ml/min. Patch clamp electrodes were pulled from borosilicate capillary tubes (1.5 mm OD, 0.86 ID, Harvard Apparatus) using a two stage puller (Narishige PP83) to a final resistance of 4-8 MΩ. The intracellular solution contained in mM: 100 CsCl, 20 TEA-Cl, 5 MgCl_2_, 2 EGTA, 10 HEPES, 5 Mg-ATP, 0.5 Li-GTP at pH 7.4. In some cases, 0.2% biocytin was included in the intracellular solution for cell labeling. Tetrodotoxin (TTX, 1 *µ*M) and tetraethylammonium (TEA, 10 mM) was added to the recording solution to block sodium and potassium currents. Whole-cell patch-clamp recordings were performed at room temperature using a Multiclamp amplifier (Molecular Devices). The signal was sampled at 20 kHz and stored for offline analysis. Spinal neurons expressing one copy of the Ca_*V*_ 1.3 gene or none were identified using a green fluorescent positive (GFP) filter in slices from newborn *HoxB8*^*Cre*^;*Cacna1d-eGFP*^+/−^ and *HoxB8*^*Cre*^;*Cacna1d-eGFP*^−/−^ mice, respectively. The amplitude and the time course of the PICs activation were estimated from a family of slow voltage-clamp bi-ramps (ascending and descending) from −80 mV to +10 mV and back to −80 mV. The linear component of the leakage current was calculated and subtracted from the traces. The ascending ramp that induces the maximum PICs was selected for calculating the PIC parameters. Changes in PICs under different conditions (control, drugs, and washout) were calculated from the PICs evoked with the same ramp. The ramp protocol was repeated three times for each condition with one minute interval in between. Cells were excluded from the study if there were clear space clamp issues (e.g. bumps and notches in the inward currents). Results are means ± SEM. Liquid junction potentials imposed by the whole-cell recordings were not corrected.

### Drug

Nimodipine (Sigma) for subcutaneous injection was used as previously reported in the literature (10 mg/kg)(46, 73). Nimodipine was dissolved in the vehicle solution (2% DMSO, 5% EtoH and physiologic solution) and kept in the dark. 20 *µ*M nifedipine was used to block L-type calcium currents *in vitro* (11). The drugs were always handled in conditions of low light.

### Immunohistochemistry

Adult mice were anaesthetized with pentobarbital, perfused with 4% (w/v) paraformaldehyde (PFA) in phosphate-buffered saline (PBS). Then, the spinal cord was dissected and post-fixed for 2 hours in 4% paraformaldehyde (PFA). Newborn mice (postnatal day 1-4) were decapitated and the spinal cord was dissected and fixed for 2 hours like for the adult tissue. After the post-fixation, the cords were rinsed in PBS, cryoprotected in 30% (wg/vol) sucrose in PBS overnight and embedded in OCT mounting medium. Transverse sections (20 *µ*m thick) were obtained on a cryostat. Sections were blocked in PBS supplemented with 5% (vol/vol) fetal bovine serum and 0.5% (wg/vol) Triton X-100 (blocking solution) before to be incubated overnight in blocking solution at 4°C with one or several of the following primary antibodies diluted in: Chicken anti-GFP (1:1000, Abcam), Goat anti-ChAT (1:100, Millipore). Secondary antibodies (Invitrogen) were incubated for 2 hours at room temperature. Fluorescent Nissl stain (NeuroTrace 647, Molecular probes) was added during the secondary antibody incubation. Slides were rinsed, mounted in Vectashield medium and scanned with a confocal microscope (Zeiss) using 10X and 20X objectives. Multiple channels were scanned sequentially to prevent fluorescence bleed.

### Cell count

For experiments in both newborn and adult mice, twelve to twenty non-adjacent cryo-sections spanning the sacral spinal cord were counted. Neurons expressing the marker of interest were manually counted using a raw Z-stack confocal image and the cell-counter plug in in imageJ. Simultaneously, five reference points were manually added to build a set of Cartesian axes with the zero centered on the central canal and the y axis parallel to the midline of the spinal cord and the x axis orthogonal to it. This way, all somas coordinates were translated and rotated to a common coordinate system. Two-dimensional Kernel density estimation was obtained using a MatLab script (DataDensityPlot) and displayed as a contour plots with the contour lines connecting points of equal density from 40% to 100% of the estimated density range with increment of 10%.

### Statistical analysis

The sample size was evaluated considering previous experiments from our laboratory and the mean and the variation of the samples. Mice were randomly allocated to the different groups for the *in vivo* experiments. The analyses were blind to the treatments (vehicle, Nimodipine). Non parametric tests were used for non-normally distributed data. All statistical tests were two-tailed. Most of the data are represented as scatterplot with bars. Bars are mean ± SEM unless specifically indicated. Significance level was indicated as n.s.=p>0.05, *=p<0.05, **=p< 0.01, ***=p<0.001.

## Supplementary Material

**Figure S1.**
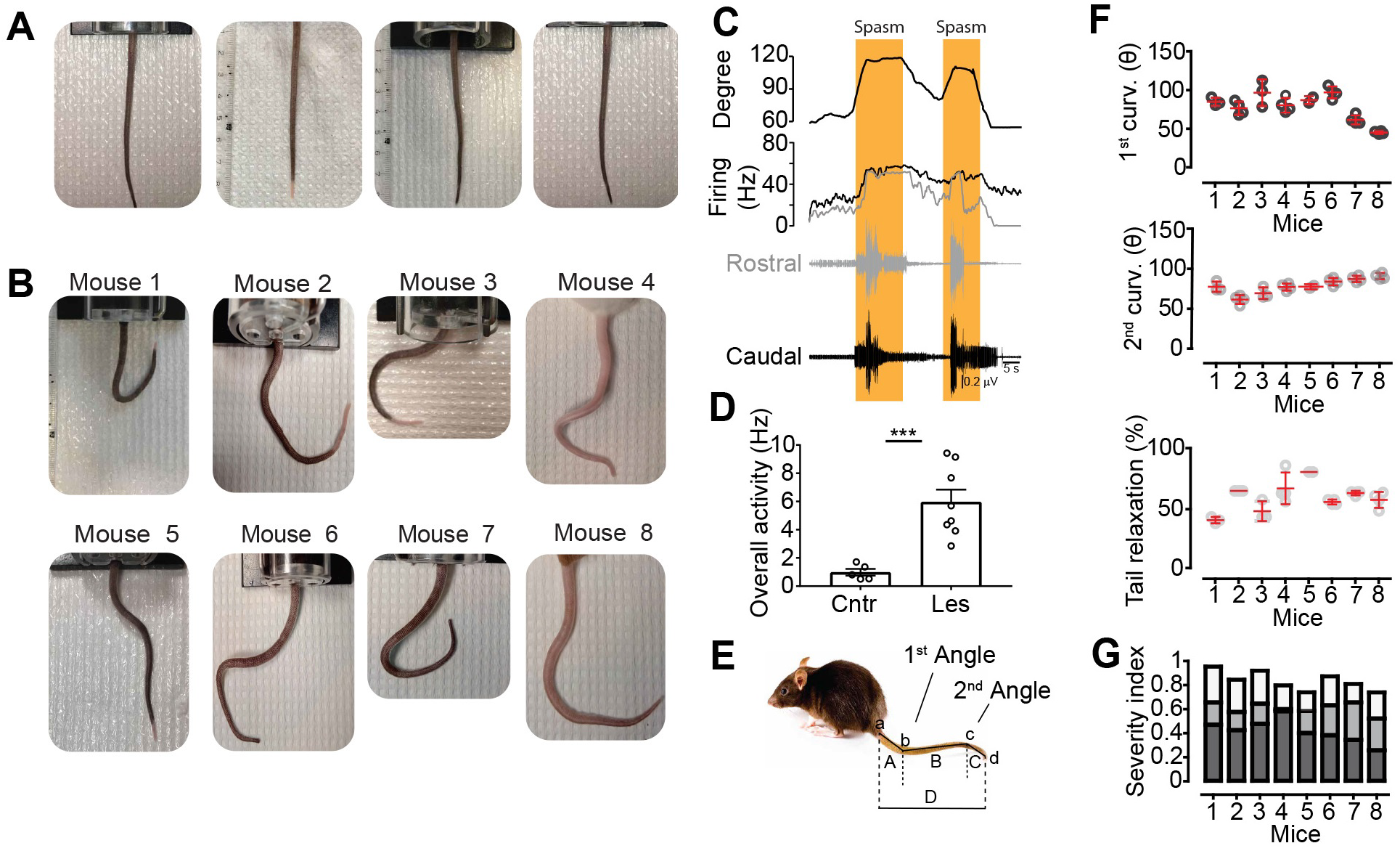
Tonic muscle contraction and muscle spasms 6 weeks after SCI.. **A-B.** Photographs of tails in wild-type unlesioned (A) and lesioned mice (B) 6 weeks after SCI. **C.** EMG recordings and firing frequency of rostral (Coccygeal 9^th^, grey) and distal (Coccygeal 17^th^, black) muscles with the simultaneous increase in tail curvature in the rostral segment (upper trace, degree) during spontaneous spasms in a lesioned wild-type mouse 6 weeks after SCI. **D.** Overall spontaneous activity (mean firing frequency) of motor units of the tail muscles from un-lesioned mice (cntr, n=131, N= 5) and lesioned mice in a (Les, n=131, N=8, unpaired Student’s t test, ***p<0.001). **E.** Schematic of the logic for tail measurements. The tail was divided in 4 points (a-d) and three segments (A-C). The segment D was generated linking the points a and d. **F.** Measurements of first curvature (upper graph), second curvature (middle graph) and relaxation (lower graph) of the tail from the mice depicted in a. **G.** Severity index calculated by weighting the first angle (darkest grey bars), second angle (dark grey bars) and relaxation (palest grey bars). See online methods for a detailed explanation.

**Figure S2.**
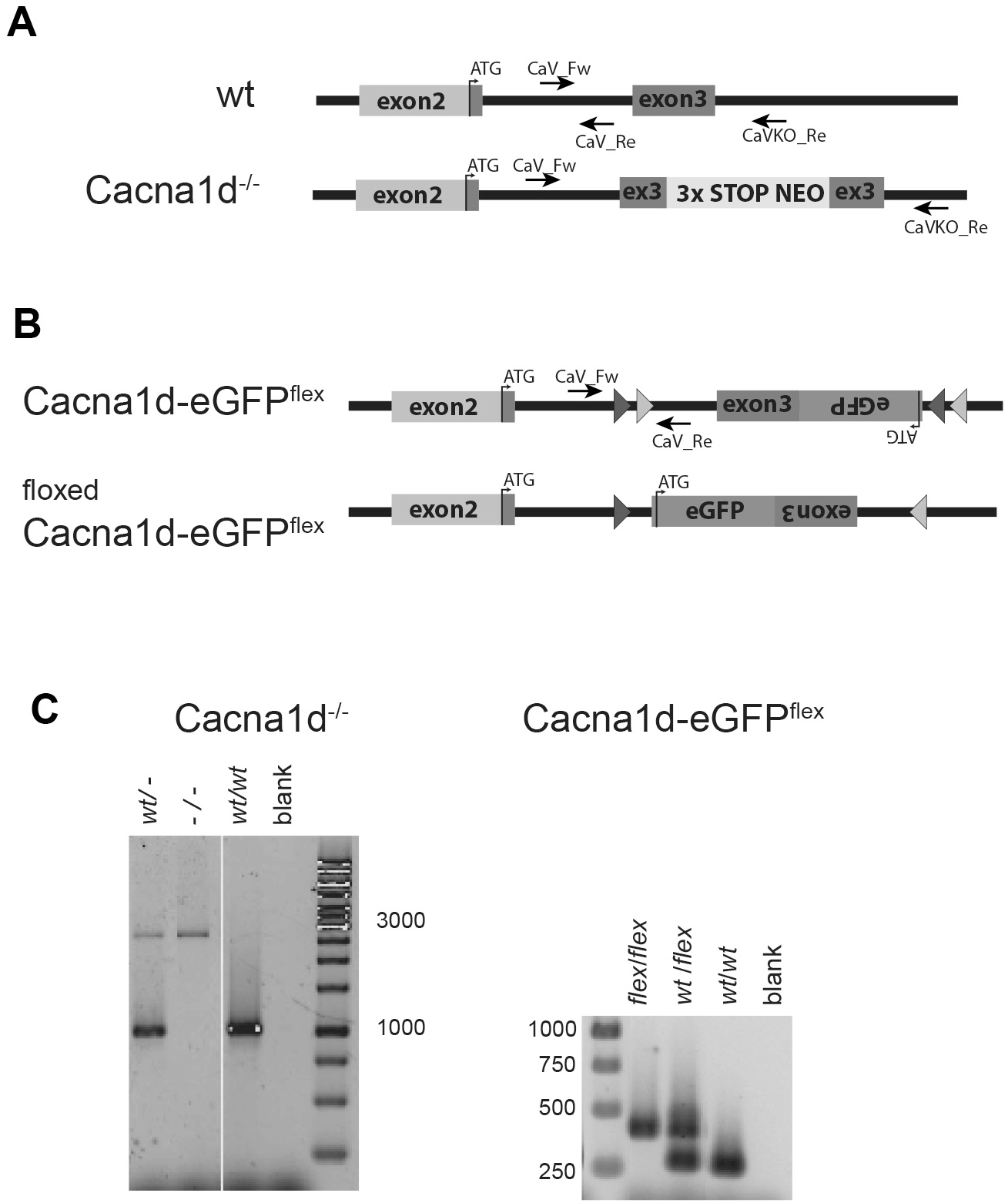
Genetics of mice used in the study. **A-B.** Schematic presentation of the two *Cacna1d* mice. The floxed version (B) is when the mouse has been subjected to Cre. For details see references 22 and 30. **C.** Typical genotyping results showing homo-, heterozygotes and wild type for the two different *Cacna1d* mice. See Material and Methods for more details.

**Figure S3.**
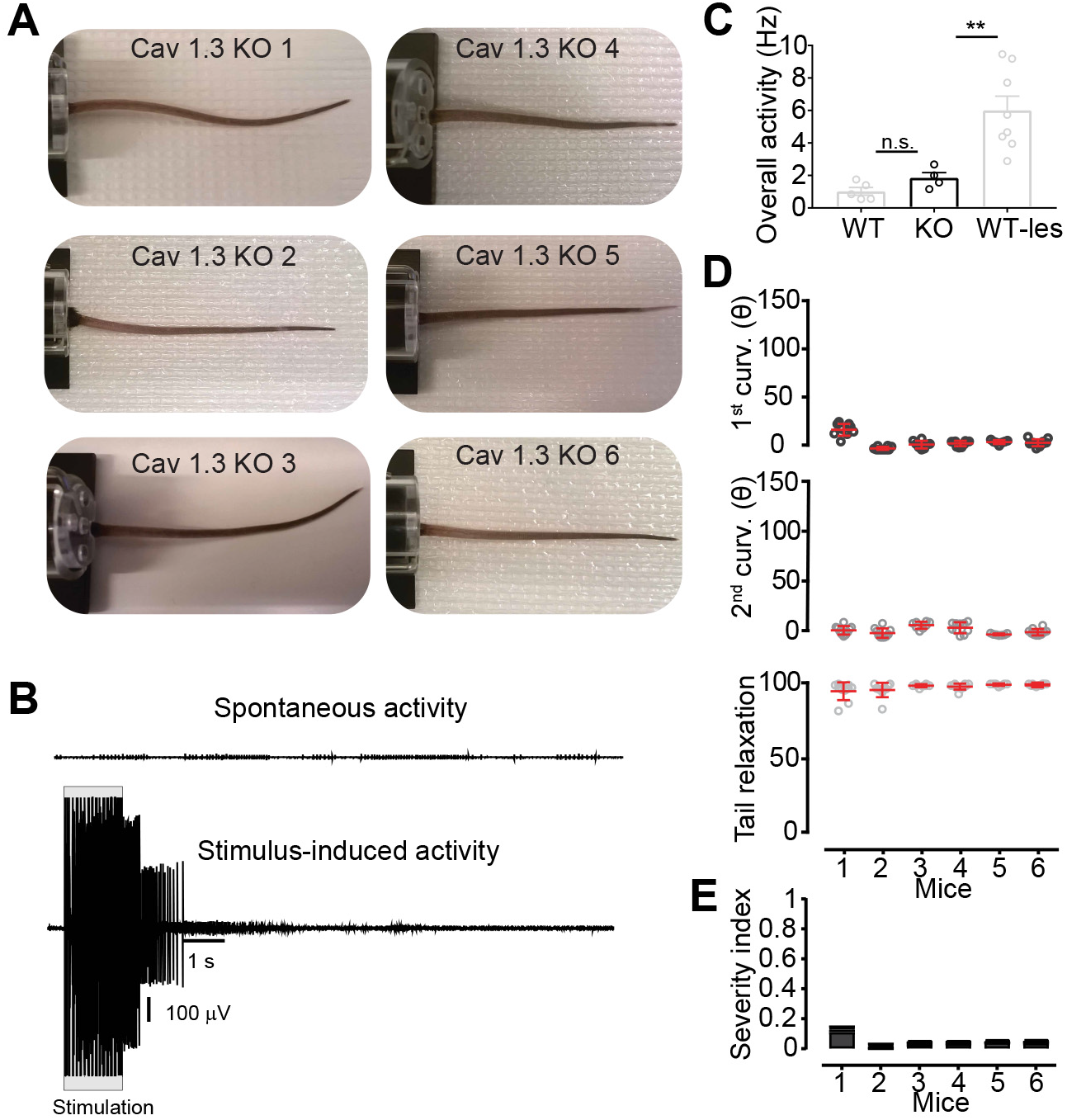
Tonic muscle contraction and spasms are not generated after SCI in mice with complete silencing of the Ca_*V*_ 1.3 calcium channels. **A.** Tails of lesioned Ca_*V*_ 1.3^−/−^ mice (Ca_*V*_ 1.3KO) six weeks after SCI. **B.** EMG recordings from the ventral tail during spontaneous activity at rest (upper trace) and stimulus-induced activity (train of electric stimuli, 500 *µ*s at 10 Hz) in a lesioned Ca_*V*_ 1.3 KO mouse. Note that large motor units can be recruited with appropriate stimulation but it does not lead to a prolonged spasm. **C.** Overall spontaneous activity from motor units (n=83) from four lesioned Ca_*V*_ 1.3 KO mice (data on wild-type and lesioned wild-type are the same as Figure S2d, One-way ANOVA, Tukey’s post-hoc, **p<0.01). **D.** Measurements of first curvature (dark grey), second curvature (light grey) and relaxation of the tail (lighter grey) for the mice depicted in a (red bars represent mean ± SEM). **E.** Severity index of Ca_*V*_ 1.3 KO mice in D.

**Figure S4.**
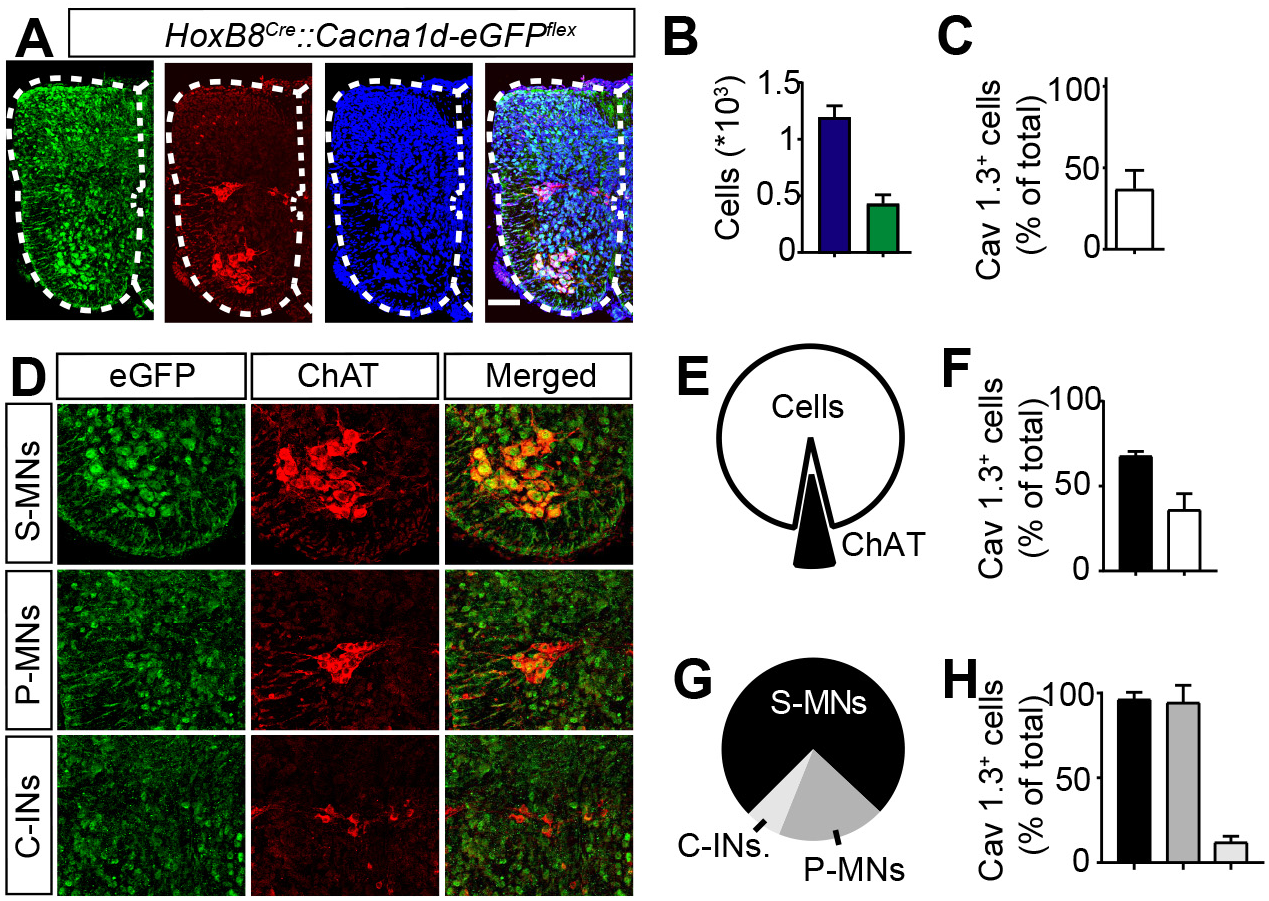
Expression of Ca_*V*_ 1.3 channels in the spinal cord of newborn mice. **A.** Confocal images of a transverse section of the spinal cord from a newborn (postnatal day one) *HoxB8*^*Cre*^;*Cacna1d-eGFP*^*Flex*^ mouse stained for eGFP (green), *ChAT* (red) and Nissl (blue). Scale bar = 100 *µ*m. **B.** Grand mean of spinal cells (Nissl, Blue) and spinal cells expressing Ca_*V*_ 1.3 channels (Ca_*V*_ 1.3^+^ cells, GFP, green) at the sacral level (n=16, N=4). **C.** Grand mean of Ca_*V*_ 1.3^+^ cells compared to the total number of spinal cells. **D.** Magnification of confocal images showing the expression of Ca_*V*_ 1.3 channels (eGFP, green) in different cholinergic neurons (ChAT^+^): somatic motor neurons (S-MNs), preganglionic motor neurons (P-MNs) and cholinergic interneurons (C-INs). **E.** Pie-chart indicating the proportion of *ChAT*extsuperscript+ / Ca_*V*_ 1.3^+^ cells of all Ca_*V*_ 1.3^+^ spinal cells. **F.** Proportion of *ChAT*^+^/Ca_*V*_ 1.3^+^ out of all ChAT^+^ neurons (black) and proportion of *ChAT*^−^/ Ca_*V*_ 1.3^+^ cells out of all nissl stained cells (white). **G.** Proportion of S-MNs (black), P-MNs (dark grey) and C-INs (pale grey) in *ChAT*^+^ neurons. **H.** Percent of S-MNs, P-MNs and C-INs expressing Ca_*V*_ 1.3 channels.

**Figure S5.**
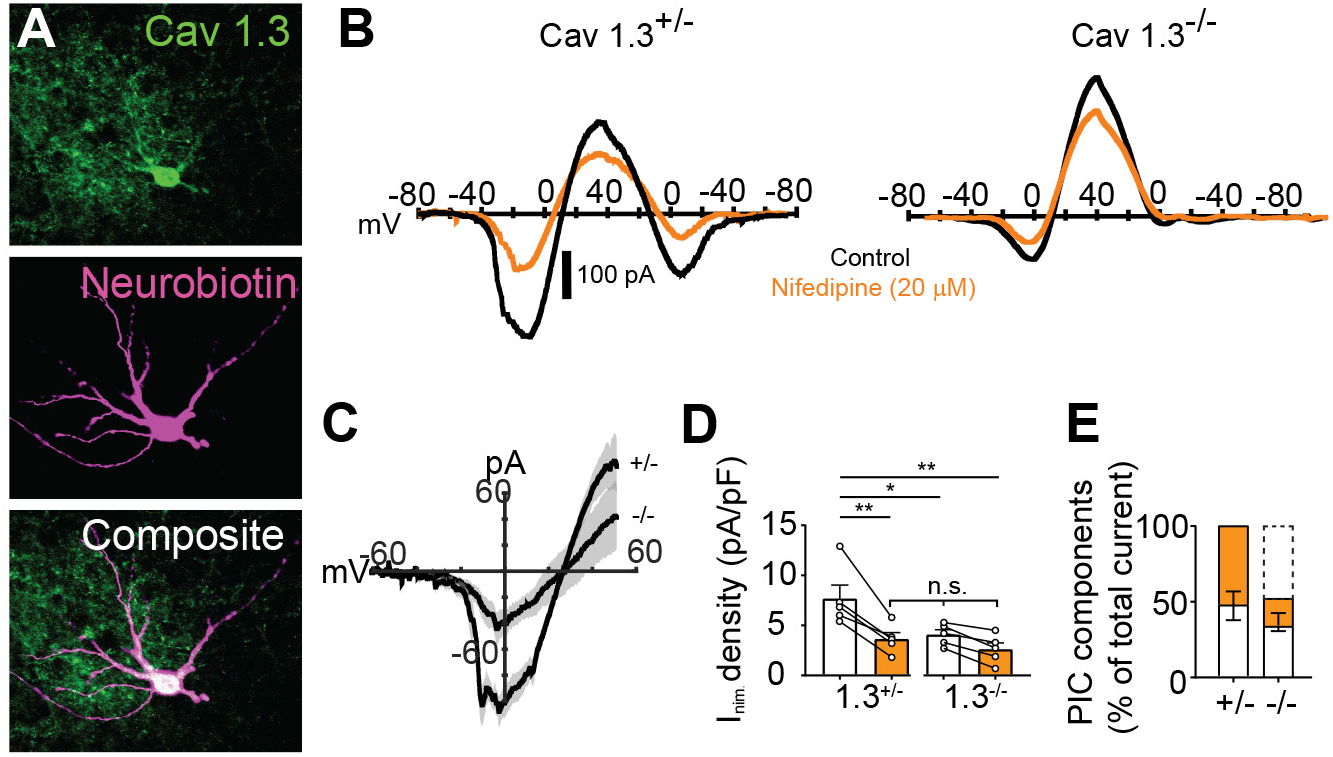
L-type calcium currents are drastically reduced in neurons of Ca_*V*_ 1.3 conditional knockout mice. **A-B.** Representative image of *HoxB8*-Ca_*V*_ 1.3^+^ neuron labeled with biotin and I-V curves in response to voltage-clamp ramps from neurons expressing one copy (Ca_*V*_ ^+/−^) or no copy of the *Cacna1d* gene (Ca_*V*_ ^−/−^) in slices from P6-8 mice spinal cords. All recordings were made in the presence of TTX 1 *µ*M, TEA 10 mM and CsCl 25 mM to block sodium and potassium currents. The persistent current (PIC) is seen as a negative slope region in the I-V curves. Black traces are control and orange traces after nifedipine (20 *µ*M). **C.** Mean current-to-voltage graph for the nifedipine-sensitive currents in neurons with on copy (+/−) or no copy (-/-) of the *Cacna1d* gene. **D.** Mean current density before and after nifedipine (±SEM, Two-way Anova, Bonferroni’s post-hoc, *p<0.05, **p<0.01). **E.** Different PIC components for all recorded cells: nifedipine sensitive PIC in orange, nifedipine insensitive in white, and Ca_*V*_ 1.3 current in white dashed line. The Ca_*V*_ current was estimated by subtraction. Data in c-e are from Ca_*V*_ ^+/−^ (n=5) and Ca_*V*_ ^−/−^ (n=5) neurons.

## Bibliography

1. C. S. Ahuja, J. R. Wilson, S. Nori, M. R. N. Kotter, C. Druschel, A. Curt, M. G. Fehlings, Traumatic spinal cord injury. Nat Rev Dis Primers 3, 17018 (2017).

2. V. Dietz, Behavior of spinal neurons deprived of supraspinal input. Nat Rev Neurol 6, 167–174 (2010).

3. S. R. Andresen, F. Biering-Sorensen, E. M. Hagen, J. F. Nielsen, F. W. Bach, N. B. Finnerup, Pain, spasticity and quality of life in individuals with traumatic spinal cord injury in Denmark. Spinal Cord 54, 973–979 (2016).

4. V. Dietz, T. Sinkjaer, Spastic movement disorder: impaired reflex function and altered muscle mechanics. Lancet Neurol 6, 725–733 (2007).

5. W. B. McKay, A. V. Ovechkin, T. W. Vitaz, D. G. Terson de Paleville, S. J. Harkema, Neurophysiological characterization of motor recovery in acute spinal cord injury. Spinal Cord 49, 421–429 (2011).

6. J. S. Krause, R. E. Carter, E. Pickelsimer, Behavioral risk factors of mortality after spinal cord injury. Arch Phys Med Rehabil 90, 95–101 (2009).

7. P. Musienko, J. Heutschi, L. Friedli, R. van den Brand, G. Courtine, Multi-system neurorehabilitative strategies to restore motor functions following severe spinal cord injury. Exp Neurol 235, 100–109 (2012).

8. F. B. Wagner, J. B. Mignardot, C. G. Le Goff-Mignardot, R. Demes-maeker, S. Komi, M. Capogrosso, A. Rowald, I. Seanez, M. Caban, E. Pirondini, M. Vat, L. A. McCracken, R. Heimgartner, I. Fodor, A. Watrin, P. Seguin, E. Paoles, K. Van Den Keybus, G. Eberle, B. Schurch, E. Pra-long, F. Becce, J. Prior, N. Buse, R. Buschman, E. Neufeld, N. Kuster, S. Carda, J. von Zitzewitz, V. Delattre, T. Denison, H. Lambert, K. Minassian, J. Bloch, G. Courtine, Targeted neurotechnology restores walking in humans with spinal cord injury. Nature 563, 65–71 (2018).

9. M. M. Adams, A. L. Hicks, Spasticity after spinal cord injury. Spinal Cord 43, 577–586 (2005).

10. J. M. Gracies, P. Nance, E. Elovic, J. McGuire, D. M. Simpson, Traditional pharmacological treatments for spasticity. Part II: General and regional treatments. Muscle Nerve Suppl 6, S92–120 (1997).

11. J. Hounsgaard, O. Kiehn, Serotonin-induced bistability of turtle motoneurones caused by a nifedipine-sensitive calcium plateau potential. J Physiol 414, 265–282 (1989).

12. Y. Li, D. J. Bennett, Persistent sodium and calcium currents cause plateau potentials in motoneurons of chronic spinal rats. J Neurophysiol 90, 857–869 (2003).

13. K. P. Carlin, K. E. Jones, Z. Jiang, L. M. Jordan, R. M. Brownstone, Dendritic L-type calcium currents in mouse spinal motoneurons: implications for bistability. Eur J Neurosci 12, 1635–1646 (2000).

14. D. J. Bennett, Y. Li, P. J. Harvey, M. Gorassini, Evidence for plateau potentials in tail motoneurons of awake chronic spinal rats with spasticity. J Neurophysiol 86, 1972–1982 (2001).

15. C. Bellardita, V. Caggiano, R. Leiras, V. Caldeira, A. Fuchs, J. Bouvier, P. Low, O. Kiehn, Spatiotemporal correlation of spinal network dynamics underlying spasms in chronic spinalized mice. Elife 6, (2017).

16. O. Roca-Lapirot, H. Radwani, F. Aby, F. Nagy, M. Landry, P. Fossat, Calcium signalling through L-type calcium channels: role in pathophysiology of spinal nociceptive transmission. Br J Pharmacol 175, 2362–2374 (2018).

17. G. W. Zamponi, J. Striessnig, A. Koschak, A. C. Dolphin, The Physiology, Pathology, and Pharmacology of Voltage-Gated Calcium Channels and Their Future Therapeutic Potential. Pharmacol Rev 67, 821–870 (2015).

18. J. T. Weber, Altered calcium signaling following traumatic brain injury. Front Pharmacol 3, 60 (2012).

19. W. Young, The role of calcium in spinal cord injury. Cent Nerv Syst Trauma 2, 109–114 (1985).

20. L. Leybaert, A. de Hemptinne, Changes of intracellular free calcium following mechanical injury in a spinal cord slice preparation. Exp Brain Res 112, 392–402 (1996).

21. R. D. Happel, K. P. Smith, N. L. Banik, J. M. Powers, E. L. Hogan, J. D. Balentine, Ca2+-accumulation in experimental spinal cord trauma. Brain Res 211, 476–479 (1981).

22. P. L. Greer, M. E. Greenberg, From synapse to nucleus: calcium-dependent gene transcription in the control of synapse development and function. Neuron 59, 846–860 (2008).

23. F. Sun, Z. He, Neuronal intrinsic barriers for axon regeneration in the adult CNS. Curr Opin Neurobiol 20, 510–518 (2010).

24. C. Bellardita, M. Marcantoni, P. Low, O. Kiehn, Sacral Spinal Cord Transection and Isolated Sacral Cord Preparation to Study Chronic Spinal Cord Injury in Adult Mice. Bio Protoc 8, e2784 (2018).

25. D. J. Bennett, M. Gorassini, K. Fouad, L. Sanelli, Y. Han, J. Cheng, Spasticity in rats with sacral spinal cord injury. J Neurotrauma 16, 69–84 (1999).

26. K. C. Murray, A. Nakae, M. J. Stephens, M. Rank, J. D’Amico, P. J. Harvey, X. Li, R. L. Harris, E. W. Ballou, R. Anelli, C. J. Heckman, T. Mashimo, R. Vavrek, L. Sanelli, M. A. Gorassini, D. J. Bennett, K. Fouad, Recovery of motoneuron and locomotor function after spinal cord injury depends on constitutive activity in 5-HT2C receptors. Nat Med 16, 694–700 (2010).

27. L. A. Ritz, R. M. Friedman, E. L. Rhoton, M. L. Sparkes, C. J. Vierck, Jr., Lesions of cat sacrocaudal spinal cord: a minimally disruptive model of injury. J Neurotrauma 9, 219–230 (1992).

28. R. L. Macdonald, T. A. Schweizer, Spontaneous subarachnoid haemorrhage. Lancet 389, 655–666 (2017).

29. M. S. Langley, E. M. Sorkin, Nimodipine. A review of its pharmaco-dynamic and pharmacokinetic properties, and therapeutic potential in cerebrovascular disease. Drugs 37, 669–699 (1989).

30. D. Tomassoni, A. Lanari, G. Silvestrelli, E. Traini, F. Amenta, Nimodipine and its use in cerebrovascular disease: evidence from recent preclinical and controlled clinical studies. Clin Exp Hypertens 30, 744–766 (2008).

31. K. J. Dougherty, S. Hochman, Spinal cord injury causes plasticity in a subpopulation of lamina I GABAergic interneurons. J Neurophysiol 100, 212–223 (2008).

32. H. Hultborn, Changes in neuronal properties and spinal reflexes during development of spasticity following spinal cord lesions and stroke: studies in animal models and patients. J Rehabil Med, 46–55 (2003).

33. J. Platzer, J. Engel, A. Schrott-Fischer, K. Stephan, S. Bova, H. Chen, H. Zheng, J. Striessnig, Congenital deafness and sinoatrial node dysfunction in mice lacking class D L-type Ca2+ channels. Cell 102, 89–97 (2000).

34. S. V. Satheesh, K. Kunert, L. Ruttiger, A. Zuccotti, K. Schonig, E. Friauf, M. Knipper, D. Bartsch, H. G. Nothwang, Retrocochlear function of the peripheral deafness gene Cacna1d. Hum Mol Genet 21, 3896–3909 (2012).

35. R. Witschi, T. Johansson, G. Morscher, L. Scheurer, J. Deschamps, H. U. Zeilhofer, Hoxb8-Cre mice: A tool for brain-sparing conditional gene deletion. Genesis 48, 596–602 (2010).

36. I. Bukreeva, G. Campi, M. Fratini, R. Spano, D. Bucci, G. Battaglia, F. Giove, A. Bravin, A. Uccelli, C. Venturi, M. Mastrogiacomo, A. Cedola, Quantitative 3D investigation of Neuronal network in mouse spinal cord model. Sci Rep 7, 41054 (2017).

37. L. Borgius, C. E. Restrepo, R. N. Leao, N. Saleh, O. Kiehn, A transgenic mouse line for molecular genetic analysis of excitatory glutamatergic neurons. Mol Cell Neurosci 45, 245–257 (2010).

38. M. Hagglund, K. J. Dougherty, L. Borgius, S. Itohara, T. Iwasato, O. Kiehn, Optogenetic dissection reveals multiple rhythmogenic modules underlying locomotion. Proc Natl Acad Sci U S A 110, 11589–11594 (2013).

39. V. Caldeira, K. J. Dougherty, L. Borgius, O. Kiehn, Spinal Hb9::Crederived excitatory interneurons contribute to rhythm generation in the mouse. Sci Rep 7, 41369 (2017).

40. Y. Dai, L. M. Jordan, Tetrodotoxin-, dihydropyridine-, and riluzole-resistant persistent inward current: novel sodium channels in rodent spinal neurons. J Neurophysiol 106, 1322–1340 (2011).

41. Y. Dai, L. M. Jordan, Multiple patterns and components of persistentinward current with serotonergic modulation in locomotor activity-related neurons in Cfos-EGFP mice. J Neurophysiol 103, 1712–1727 (2010).

42. K. P. Carlin, Z. Jiang, R. M. Brownstone, Characterization of calcium currents in functionally mature mouse spinal motoneurons. Eur J Neurosci 12, 1624–1634 (2000).

43. G. S. Bhumbra, M. Beato, Recurrent excitation between motoneurones propagates across segments and is purely glutamatergic. PLoS Biol 16, e2003586 (2018).

44. H. Nishimaru, C. E. Restrepo, J. Ryge, Y. Yanagawa, O. Kiehn, Mammalian motor neurons corelease glutamate and acetylcholine at central synapses. Proc Natl Acad Sci U S A 102, 5245–5249 (2005).

45. J. M. Cregg, K. A. Chu, L. E. Hager, R. S. J. Maggard, D. R. Stoltz, M. Edmond, W. J. Alilain, P. Philippidou, L. T. Landmesser, J. Silver, A Latent Propriospinal Network Can Restore Diaphragm Function after High Cervical Spinal Cord Injury. Cell Rep 21, 654–665 (2017).

46. A. Schampel, O. Volovitch, T. Koeniger, C. J. Scholz, S. Jorg, R. A. Linker, E. Wischmeyer, M. Wunsch, J. W. Hell, S. Ergun, S. Kuerten, Nimodipine fosters remyelination in a mouse model of multiple sclerosis and induces microglia-specific apoptosis. Proc Natl Acad Sci U S A 114, E3295–E3304 (2017).

47. X. Wang, N. Lou, Q. Xu, G. F. Tian, W. G. Peng, X. Han, J. Kang, T. Takano, M. Nedergaard, Astrocytic Ca2+ signaling evoked by sensory stimulation in vivo. Nat Neurosci 9, 816–823 (2006).

48. C. Skold, R. Levi, A. Seiger, Spasticity after traumatic spinal cord injury: nature, severity, and location. Arch Phys Med Rehabil 80, 1548–1557 (1999).

49. K. A. Holtz, R. Lipson, V. K. Noonan, B. K. Kwon, P. B. Mills, Prevalence and Effect of Problematic Spasticity After Traumatic Spinal Cord Injury. Arch Phys Med Rehabil 98, 1132–1138 (2017).

50. E. Chang, N. Ghosh, D. Yanni, S. Lee, D. Alexandru, T. Mozaffar, A Review of Spasticity Treatments: Pharmacological and Interventional Approaches. Crit Rev Phys Rehabil Med 25, 11–22 (2013).

51. J. N. Guzman, J. Sanchez-Padilla, D. Wokosin, J. Kondapalli, E. Ilijic, P. T. Schumacker, D. J. Surmeier, Oxidant stress evoked by pacemaking in dopaminergic neurons is attenuated by DJ-1. Nature 468, 696–700 (2010).

52. F. M. Warner, J. J. Cragg, C. R. Jutzeler, F. Rohrich, N. Weidner, M. Saur, D. D. Maier, C. Schuld, A. Curt, J. K. Kramer, Early Administration of Gabapentinoids Improves Motor Recovery after Human Spinal Cord Injury. Cell Rep 18, 1614–1618 (2017).

53. A. Tedeschi, S. Dupraz, C. J. Laskowski, J. Xue, T. Ulas, M. Beyer, J. L. Schultze, F. Bradke, The Calcium Channel Subunit Alpha2delta2 Suppresses Axon Regeneration in the Adult CNS. Neuron 92, 419–434 (2016).

54. V. Dietz, M. E. Schwab, From the Rodent Spinal Cord Injury Model to Human Application: Promises and Challenges. J Neurotrauma 34, 1826–1830 (2017).

55. J. Hounsgaard, O. Kiehn, Ca++ dependent bistability induced by serotonin in spinal motoneurons. Exp Brain Res 57, 422–425 (1985).

56. C. J. Heckmann, M. A. Gorassini, D. J. Bennett, Persistent inward currents in motoneuron dendrites: implications for motor output. Muscle Nerve 31, 135–156 (2005).

57. U. I. Scholl, G. Goh, G. Stolting, R. C. de Oliveira, M. Choi, J. D. Overton, A. L. Fonseca, R. Korah, L. F. Starker, J. W. Kunstman, M. L. Prasad, E. A. Hartung, N. Mauras, M. R. Benson, T. Brady, J. R. Shapiro, E. Loring, C. Nelson-Williams, S. K. Libutti, S. Mane, P. Hellman, G. Westin, G. Akerstrom, P. Bjorklund, T. Carling, C. Fahlke, P. Hidalgo, R. P. Lifton, Somatic and germline CACNA1D calcium channel mutations in aldosterone-producing adenomas and primary aldosteronism. Nat Genet 45, 1050–1054 (2013).

58. A. Andrade, A. Sandoval, R. Gonzalez-Ramirez, D. Lipscombe, K. P. Campbell, R. Felix, The alpha(2)delta subunit augments functional expression and modifies the pharmacology of Ca(V)1.3 L-type channels. Cell Calcium 46, 282–292 (2009).

59. A. Pinggera, J. Striessnig, Cav 1.3 (CACNA1D) L-type Ca(2+) channel dysfunction in CNS disorders. J Physiol 594, 5839–5849 (2016).

60. C. M. Moreno, R. E. Dixon, S. Tajada, C. Yuan, X. Opitz-Araya, M. D. Binder, L. F. Santana, Ca(2+) entry into neurons is facilitated by cooperative gating of clustered CaV1.3 channels. Elife 5, (2016).

61. O. Vivas, C. M. Moreno, L. F. Santana, B. Hille, Proximal clustering between BK and CaV1.3 channels promotes functional coupling and BK channel activation at low voltage. Elife 6, (2017).

62. C. Brocard, V. Plantier, P. Boulenguez, S. Liabeuf, M. Bouhadfane, A. Viallat-Lieutaud, L. Vinay, F. Brocard, Cleavage of Na(+) channels by calpain increases persistent Na(+) current and promotes spasticity after spinal cord injury. Nat Med 22, 404–411 (2016).

63. J. Niu, I. E. Dick, W. Yang, M. A. Bamgboye, D. T. Yue, G. Tomaselli, T. Inoue, M. Ben-Johny, Allosteric regulators selectively prevent Ca(2+)- feedback of CaV and NaV channels. Elife 7, (2018).

64. M. L. McEwen, P. G. Sullivan, A. G. Rabchevsky, J. E. Springer, Targeting mitochondrial function for the treatment of acute spinal cord injury. Neurotherapeutics 8, 168–179 (2011).

65. M. E. Schwab, D. Bartholdi, Degeneration and regeneration of axons in the lesioned spinal cord. Physiol Rev 76, 319–370 (1996).

66. G. Mandolesi, F. Madeddu, Y. Bozzi, L. Maffei, G. M. Ratto, Acute physiological response of mammalian central neurons to axotomy: ionic regulation and electrical activity. FASEB J 18, 1934–1936 (2004).

67. A. Tedeschi, F. Bradke, Spatial and temporal arrangement of neuronal intrinsic and extrinsic mechanisms controlling axon regeneration. Curr Opin Neurobiol 42, 118–127 (2017).

68. Y. Li, A. M. Lucas-Osma, S. Black, M. V. Bandet, M. J. Stephens, R. Vavrek, L. Sanelli, K. K. Fenrich, A. F. Di Narzo, S. Dracheva, I. R. Winship, K. Fouad, D. J. Bennett, Pericytes impair capillary blood flow and motor function after chronic spinal cord injury. Nat Med 23, 733–741 (2017).

69. S. K. Agrawal, R. Nashmi, M. G. Fehlings, Role of L- and N-type calcium channels in the pathophysiology of traumatic spinal cord white matter injury. Neuroscience 99, 179–188 (2000).

70. T. Winkler, H. S. Sharma, E. Stalberg, R. D. Badgaiyan, T. Gordh, J. Westman, An L-type calcium channel blocker, nimodipine influences trauma induced spinal cord conduction and axonal injury in the rat. Acta Neurochir Suppl 86, 425–432 (2003).

71. M. A. Anderson, T. M. O’Shea, J. E. Burda, Y. Ao, S. L. Barlatey, A. M. Bernstein, J. H. Kim, N. D. James, A. Rogers, B. Kato, A. L. Wollenberg, R. Kawaguchi, G. Coppola, C. Wang, T. J. Deming, Z. He, G. Courtine, M. V. Sofroniew, Required growth facilitators propel axon regeneration across complete spinal cord injury. Nature 561, 396–400 (2018).

72. J. M. Cregg, M. A. DePaul, A. R. Filous, B. T. Lang, A. Tran, J. Silver, Functional regeneration beyond the glial scar. Exp Neurol 253, 197–207 (2014).

73. A. Singh, P. Verma, G. Balaji, S. Samantaray, K. P. Mohanakumar, Ni-modipine, an L-type calcium channel blocker attenuates mitochondrial dys-functions to protect against 1-methyl-4-phenyl-1,2,3,6-tetrahydropyridine-induced Parkinsonism in mice. Neurochem Int 99, 221–232 (2016).

74. D. J. Bennett, M. Gorassini, K. Fouad, L. Sanelli, Y. Han, J. Cheng, Spasticity in rats with sacral spinal cord injury. J Neurotrauma 16, 69–84 (1999).

75. J. H. Lim, L. Boozer, C. L. Mariani, J. A. Piedrahita, N. J. Olby, Generation and characterization of neurospheres from canine adipose tissue-derived stromal cells. Cell Reprogram 12, 417–425 (2010).

76. L. A. Ritz, R. M. Friedman, E. L. Rhoton, M. L. Sparkes, C. J. Vierck, Jr., Lesions of cat sacrocaudal spinal cord: a minimally disruptive model of injury. J Neurotrauma 9, 219–230 (1992).

